# Local IFNγ signaling contributes to the regenerative decline of aged alveolar progenitor cells

**DOI:** 10.64898/2026.04.07.716929

**Authors:** Jake Jensen, Kai Guo, Janine Gote-Schniering, Meeta Mistry, Zane Orinska, Jia-qi Wang, Maria Camila Melo-Narvaez, Gowtham Boosarpu, Amine Chahin, Margherita Paschini, Madison Seymour, Patrizia Pessina, Susanna M. Dang, Qianjiang Hu, Shannan Ho Sui, Melanie Königshoff, Mareike Lehmann, Silke Meiners, Carla F. Kim

**Affiliations:** Department of Genetics, Harvard Medical School; Boston, MA, USA; Stem Cell and Regenerative Biology Program, Department of Pediatrics, Division of Hematology/Oncology and Pulmonary Medicine, Boston Children’s Hospital; Boston, MA, USA; Institute of Experimental Medicine, Christian Albrecht University of Kiel; Kiel, Germany; Research Center Borstel, Leibniz Lung Center, Airway Research Center North (ARCN), Member of the German Center for Lung Research (DZL), Borstel, Germany; Harvard Chan Bioinformatics Core, Harvard T.H. Chan School of Public Health, Boston, MA, USA; Department of Rheumatology and Immunology, Department of Pulmonary Medicine, Allergology and Clinical Immunology, Inselspital, Bern University Hospital, University of Bern, Bern, Switzerland; Lung Precision Medicine (LPM), Department for BioMedical Research (DBMR), University of Bern, Bern, Switzerland; Center of Lung Aging and Regeneration, Division of Pulmonary, Allergy, Critical Care, and Sleep Medicine, University of Pittsburgh School of Medicine, Pittsburgh, PA, USA; Institute for Lung Research, Philipps-University Marburg, Member of the German Center for Lung Research (DZL), Marburg, Germany; Comprehensive Pneumology Center (CPC), Institute of Lung Health and Immunity, Helmholtz Zentrum München, Member of the German Center for Lung Research (DZL), Munich, Germany; Institute for Lung Health (ILH), Giessen, Germany; Harvard Stem Cell Institute, Cambridge, MA, USA

**Author notes:** These authors contributed equally to this work (Jake Jensen, Kai Guo, Janine Gote-Schniering).

## Abstract

The lungs are highly susceptible to chronic disease in advanced age, likely due to the uniquely compromised repair function of alveolar type II (AT2) cells, facultative progenitor cells that maintain the gas exchange surface. Using aging mouse models, single-cell sequencing, and *ex vivo* organoid assays, we found that homeostatic aged AT2 cells exhibited an Interferon γ (IFNγ) response associated with IFNγ+ CD8+ T cells in tertiary lymphoid structures (TLS). Aged AT2 cells exhibit impaired regeneration in organoid assays and lost markers of an IFNγ response outside the lung microenvironment, demonstrating that elevated local IFNγ influences the state of AT2 cells. Neutralization of IFNγ signaling and immunoproteasome knockout mice with attenuated IFNγ levels partially rescued aged AT2 cell regeneration. Our findings demonstrate that local IFN*γ* signaling in aging lungs actively represses alveolar regeneration, establishing chronic inflammatory signaling as a cause of age-related decline in the lung. Halting chronic inflammatory processes restored alveolar regeneration and may provide a means to improve lung health in old age.

## Introduction

Old age is well-understood as one of the largest risk factors for respiratory disease incidence and severity (*1*), yet how aging impacts the regenerative function of lung progenitor cells is poorly understood. Conditions that emerge from lung disrepair, such as Chronic Obstructive Pulmonary Disease (COPD) which accounts for 5% of worldwide mortality (*2*), represent an enormous public health burden with susceptibility in older individuals. The recently described age-related decline in alveolar type II (AT2) cell number and progenitor function almost certainly plays a role in poor age-related lung disease outcomes (*3–7*). Although usually quiescent, AT2 cells act as the key reparative cell of the gas-exchange surface by engaging a regenerative program to renew lost AT2 cells and differentiate into alveolar type I (AT1) cells during injury or turnover (*8*). Decline in AT2 regenerative function is seen in the reduced AT2 proliferative response after alveolar injury in aged lungs (*3–7*), the reduced capacity of aged AT2 cells to form organoids in culture (*3,5–6*), and the inability to regenerate alveoli after pneumonectomy (*9*). This contrasts with airway cell populations, such as the secretory cells and multipotent bronchioalveolar stem cells, which maintain organoid formation capacity and a strong proliferative response to injury in aged animals (*3*). Cell-intrinsic metabolic factors that contribute to AT2 cell decline in old age are now emerging, including the disruption of glycolysis and iron homeostasis (*4,5*), but there is little understanding of the influence of the aging microenvironment.

AT2 cells experience a variety of unique inflammatory signals in the aged lung (*10*), which are now being newly appreciated for their role in lung regeneration, such as IL-1β (*11,12*), TNFα (*12*), downstream NF-κB (*12,13*), and interferon (IFN) signaling (*14–17*). IFN signaling has a wide range of differential impacts on cell growth depending on the specific IFN ligand (IFNα, IFNβ, IFNγ, IFNλ, etc.), signaling subtype (type I, II, or III), organ, and cell type (*18*). IFN is well known to reprogram epithelial cells during infection or cancer by initiating heightened interactions with surveilling CD8+ T cells via upregulation of MHC class I complex and the immunoproteasome, which breaks down cellular proteins into peptide antigens for presentation by MHC-I on the cell surface to CD8+ T cells (*18–20*). Despite prior observations of elevated IFN in the aged lung (*21,22*) and strong historical literature on IFN signaling (*18*), its role has never been studied in the aged lung. Here we sought to decipher the source of elevated interferon signaling in the aged lung and what role it may play in the declining progenitor function of aged AT2 cells.

## Results

### Aged AT2 cells exhibit a conserved inflammatory IFNγ-response signature dependent on the aged lung microenvironment

To determine which transcriptional changes are conserved despite stochastic aging processes, we performed bulk RNA-seq of FACS-isolated young (2 months) and old (18 months) AT2 cells (Fig. 1A, Supp Fig. 1A), then compared differentially expressed genes to two published murine aged lung transcriptomic datasets with AT2 cells (Fig. 1B). Common among all the differentially upregulated genes in each dataset’s aged AT2 cells were genes crucial for the MHC-I antigen presentation process, including subunits of the classical MHC-I complex (*B2m, H2-K1, H2-D1*) and the immunoproteasome catalytic subunits *Psmb8* and *Psmb9*. Upregulation of the immunoproteasome in aged AT2 cells matches our previous work showing elevated immunoproteasome subunit expression in aged whole lung (*53*). scRNA sequencing from the Human lung cell atlas (*23*) was further explored to determine if an MHC-I complex and immunoproteasome signature was observed in aged human AT2 cells. Aged human AT2 cells had significantly elevated gene signature scores for MHC-I complex and immunoproteasome function compared to young cells (Fig. 1C), reflective of shared transcriptional changes with murine aged AT2 cells.

**Figure 1.**
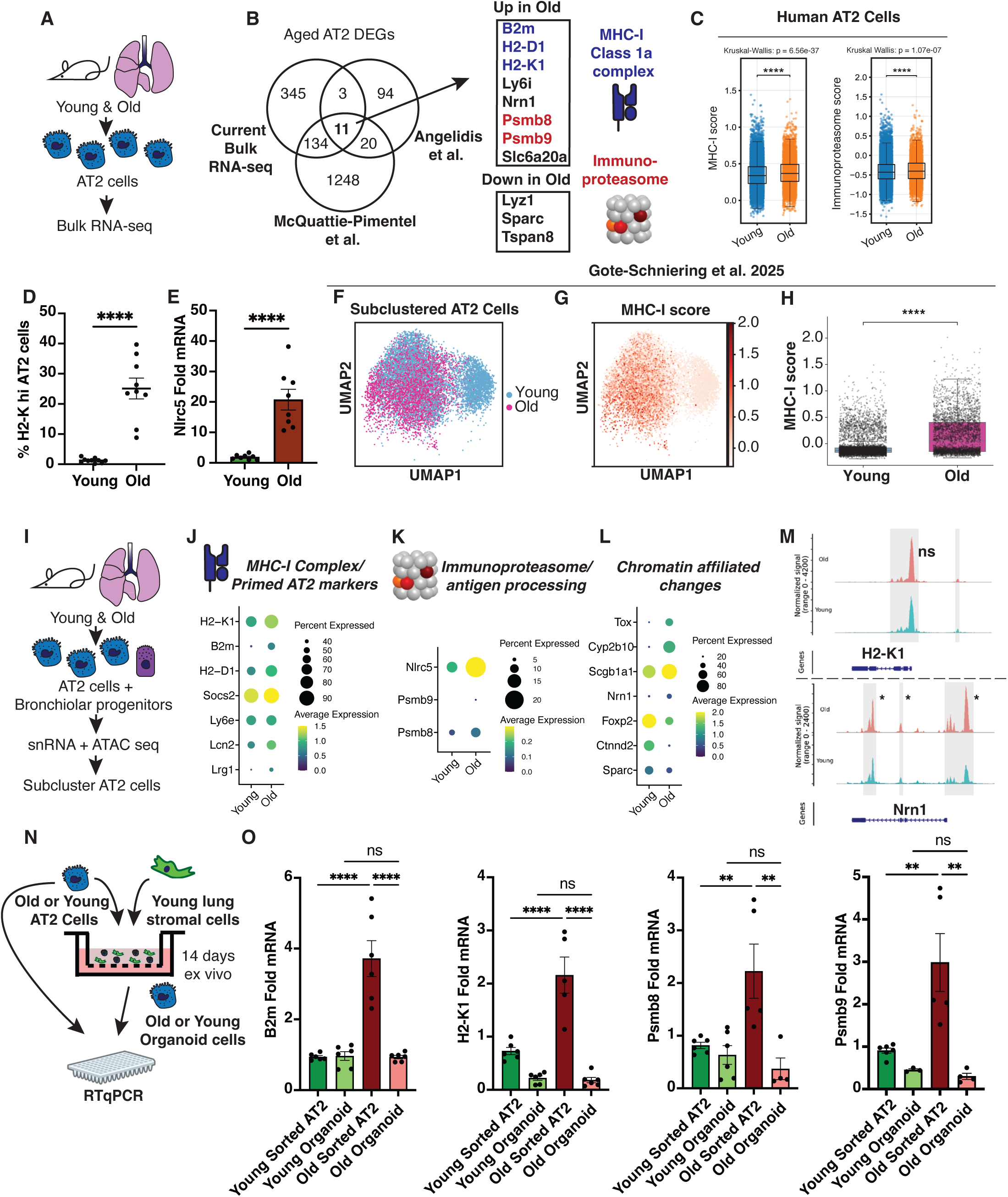
Aged AT2 cells exhibit a conserved inflammatory IFNγ-response signature dependent on the aged lung microenvironment. A) AT2 cells were isolated from aged mice (18 months) and young controls (2 months) for bulk RNA sequencing. B) Significantly differentially expressed genes in aged AT2 cells (adjusted p-value <0.05, log fold change >0.3) were compared with other published aged AT2 cell transcriptomic datasets, including Angelidis et al. 2019 (comparing 24 months old and 3 month-old C57BL/6 mice, *19*) and McQuattie-Pimental et al. 2021 (comparing 18 months old and 4 month old C57BL/6 mice, *53*). Shared gene expression changes across these datasets with advanced age are shown. C) Human lung cell atlas (*23*) comparison of MHC-I complex and immunoproteasome genes in AT2 cells across age cohorts; young (<30 years), old (>60 years). D) Flow cytometry of anti-H2-K_b_ surface staining in young and old murine AT2 cells (CD31-CD45-EPCAM+Sca1-CD24-) to measure surface MHC-I complex. E) RTqPCR of Nlrc5 mRNA from sorted young and old AT2 cells. F) UMAP representation of AT2 cells from young and aged uninjured mouse lungs (reanalysis of Gote-Schniering et al. *26*). G) UMAP representation displaying expression of MHC-I complex gene scores within young and aged murine AT2 cells H) Boxplots displaying single-cell resolved MHC-I complex gene scores between young and aged AT2 cells. I) AT2 cells and bronchiolar progenitor were isolated from aged mice (24 months) and young controls (2 months) for single nuclear multiomic sequencing (joint RNA and ATAC sequencing). In subclustered AT2 cells (see Supp Fig. 1F-G for annotation) key genes of the MHC-I complex and activated AT2 state (J), immunoproteasome (K), and select chromatin-associated gene expression changes (L) were compared between young and old AT2 cells by dotplot from a total of 285 differentially expressed genes with >0.25 fold change. M) snATAC-seq coverage plots visualizing reads around the H2-K1 and Nrn1 gene as a measure of chromatin accessibility in young and old AT2 cells. Gray boxes represent peaks called by MACS2 and significantly linked peak changes marked. N) Isolation of aged and young AT2 cells and matching *ex vivo* organoid Epcam+ cells after 14 days in culture with young lung stromal cells. O) RTqPCR comparing MHC-I complex and immunoproteasome gene expression in freshly sorted (*in vivo)* and cultured (*ex vivo*) aged AT2 cells. *P < 0.05, **P < 0.01, ***P < 0.001, ****P < 0.0001.

Though homeostatic AT2 cells express low levels of MHC-I complex as is common in slow-growing progenitors (*24*), flow cytometry of AT2 cells from aged mice showed significantly elevated MHC-I complex surface expression (Fig. 1D, Supp Fig. 1B). Additionally, RTqPCR of AT2 cells sorted from aged mice showed a near 20-fold increase in expression of *Nlrc5* (Fig. 1E), a transcriptional activator of MHC-I-mediated antigen presentation and processing machinery known to be responsive to IFN signaling (*25*). To further explore differences in AT2 cells with elevated MHC-I, we stratified AT2 cells from our previously generated aged mouse lung scRNA sequencing dataset (*26*) by MHC-I score and compared the top 25% MHC-I high AT2 with the remaining AT2 cells (Fig. 1F-G, Supp Fig. 1C-E). These MHC-I high AT2 cells had significantly reduced gene expression of G2-M checkpoint and mitotic spindle proteins and elevated stress signaling such as ROS (Supp Fig. 1E), highlighting a plausible link to reduced self-renewal capacity and the proliferative defects seen in aged AT2 cells (*3–7*).

To investigate gene regulatory mechanisms driving conserved transcriptional changes in aged AT2 cells, we enriched for AT2 cells and bronchiolar progenitors via sorting from young and old mouse lungs (2 and 24 months old, respectively) for single-nucleus joint RNA and ATAC sequencing (snMultiomic-seq, Fig. 1I). After cell annotation based on a joint RNA and ATAC clustering analysis, AT2 cells comprised the majority of the dataset (Supp Fig. 1F-G), as expected. Subclustered AT2 cells separated by age in individual RNA and ATAC UMAP projections (Supp. Fig. 1H-I). MHC-I and immunoproteasome mRNA was elevated in aged AT2 cells as expected (Fig. 1J-K). Gene set enrichment analysis (GSEA) of differentially expressed genes identified elevated interferon gamma (IFNγ) response genes and various metabolic pathways as elevated pathways in aged AT2 cells, with terms related to MHC-I-mediated antigen presentation as seen across aged AT2 datasets (Supp Fig. 1J). In contrast, snATAC sequencing did not identify age-related differences in chromatin accessibility at MHC-I-mediated antigen presentation genes, although other aged AT2 cell marker genes (*Nrn1, Scgb1a1, Cyp2b10, Tox*) exhibited differential chromatin accessibility linked to gene expression changes (Fig. 1L-M). The regulatory regions of *B2m, H2-K1, Psmb8,* and *Psmb9* had unaltered chromatin accessibility in aged AT2 by snATAC-seq (Fig. 1M, Data S1), providing evidence that cell-intrinsic chromatin regulation is not responsible for the observed IFN-responsive state. Aged AT2 cells also highly expressed *Lrg1* and *Lcn2*, markers of a previously described activated or primed AT2 cell state (*11,27*), but no elevation of the *Krt8*^hi^ AT2-AT1 transitional state implicated in several lung pathologies (Fig. 1J, Supp Fig. 1K, Data S2, *27-31*).

We next sought to test whether the aged lung microenvironment is necessary to perpetuate the IFN-responsive state of aged AT2 cells. AT2 cell aging markers were measured by RTqPCR in sorted AT2 cells from young and old mice and then compared to matched alveolar organoids derived from the same AT2 cells, grown similarly to previous studies (Fig. 1NN) (*14, 29, 32*). IFN-responsive AT2 cell aging markers, such as components of classical MHC-I complex (*B2m, H2-K1*) and the immunoproteasome (*Psmb8, Psmb9*), were elevated in freshly isolated aged AT2 cells compared to young cells as expected (Fig. 1O). However, expression of these genes was no longer elevated in lung stroma-supported or feeder-free organoids derived from aged AT2 cells compared to young cells (Fig. 1O, Supp Fig. 2A-C). In contrast, the conserved aged AT2 marker gene *Nrn1* that was linked with elevated chromatin accessibility in aged AT2 cells (Fig. 1N) remained overexpressed in alveolar organoids derived from aged cells (Supp Fig. 2E-F). This suggested that the aged lung microenvironment is required to maintain the IFN-responsive state of aged AT2 cells.

**Figure 2.**
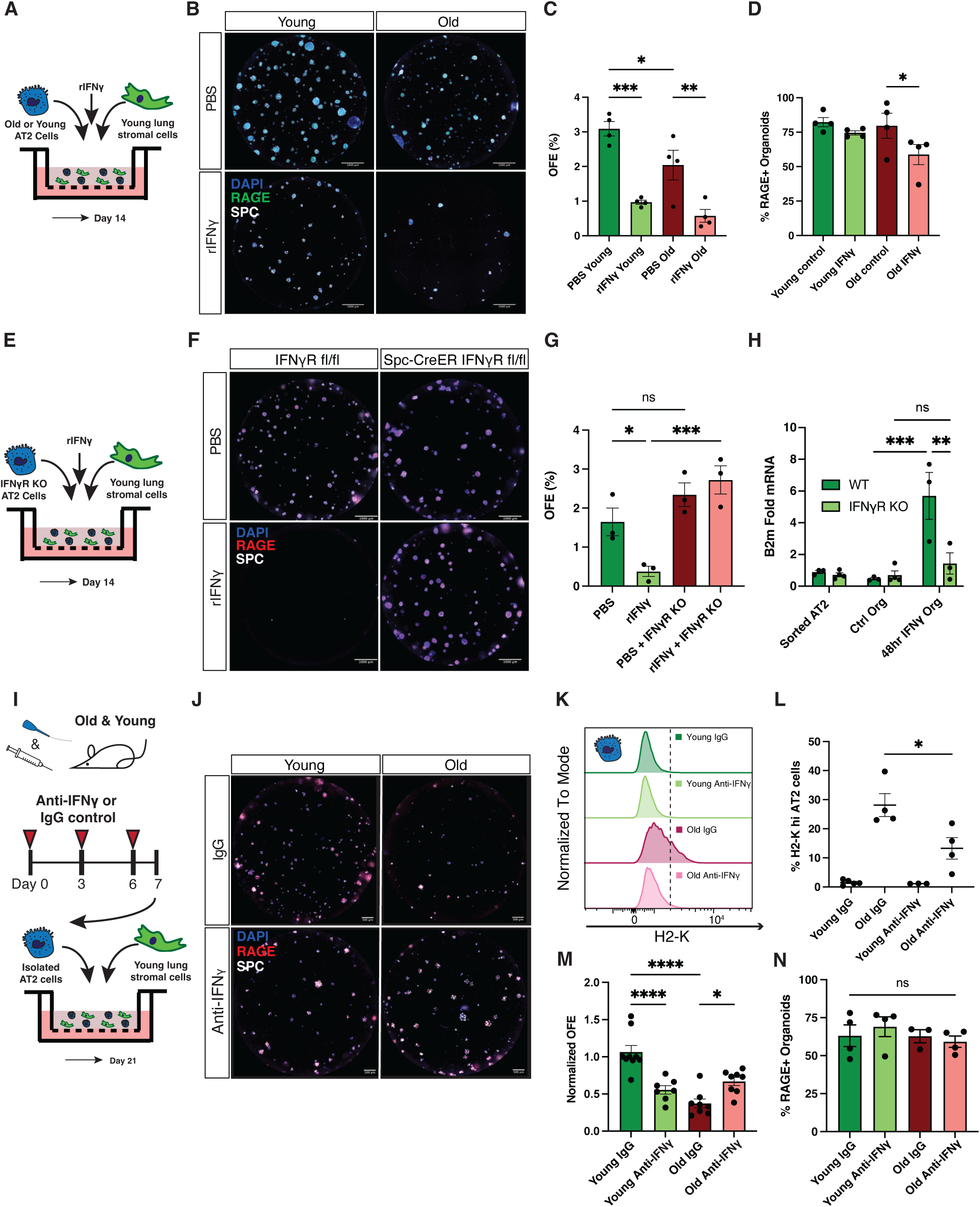
Local IFNγ signaling represses AT2 cell regeneration in the aged lung and *ex vivo* organoid assays. A) Experiment diagram of alveolar organoid cultures derived from FACS isolated young or old AT2 cells with added rIFNγ (10ng/μl) or PBS vehicle control. B) Representative well images of AT2 (Sftpc+) and AT1 (RAGE+) cell immunofluorescence, (C) quantification of organoid forming efficiency, and (D) quantification of Sftpc+ organoids also containing RAGE+ AT1 cells by immunofluorescence. E) Experimental diagram of alveolar organoid cultures derived from FACS isolated IFNγR1 conditional knockout or control AT2 cells with added rIFNγ (10ng/μl) or PBS vehicle control. F) Representative well images of AT2 (Sftpc+) and AT1 (RAGE+) cell immunofluorescence and (G) quantification of organoid forming efficiency. H) RTqPCR of Epcam+ alveolar organoid cells from IFNγR1 KO and control mice after 14 days in culture with 48 hour exposure to rIFNγ treatment (10ng/mL). I) Experimental diagram of anti-IFNγ treatment in young or old mice follow by FACS and *ex vivo* organoid assays, with (J) immunofluorescent imaging of representative wells with anti-Sftpc AT2 cell marker (white) and anti-RAGE AT1 cell marker (red). K) Representative histograms of surface MHC-I complex component H2-K_b_ across age and with IFNγ neutralization in flow cytometry of AT2 cells, including (L) quantification of H2-K_b_ high cells. M) Quantification of organoid forming efficiency as proxy for AT2 self-renewal and (F) ratio of RAGE+ alveolar organoids as a proxy for AT1 differentiation. *P < 0.05, **P < 0.01, ***P < 0.00, ****P < 0.0001.

We asked which of the many forms of interferon signaling could explain the elevated antigen presentation signature of aged AT2 cells. We reasoned that a type II IFNγ response may be dominant in aged AT2 cells compared to type I or type III interferon, due to its elevated levels in the aged lung (*21,22*), the dependence on IFNγR for elevated MHC-I expression in AT2 cells (*33*), and the specific gene expression profile of elevated *Nlrc5, Socs2*, and antigen presentation machinery (Fig. 1J-L, *33-35*). Whereas these genes are associated with transcriptional control by all types of IFN signaling, no canonical ISGF3-dependent genes (*Isg15, Ifit1, Oas3*, etc.), which are more specific to type I or type III interferon signaling (*18, 35*), were found to be elevated. This supported our subsequent investigation into the role of IFNγ in aged lung.

### Local IFNγ signaling represses AT2 cell regeneration in the aged lung and *ex vivo* organoids

To understand whether local IFNγ signaling was sufficient to disrupt AT2 cell regeneration, we treated alveolar organoid cultures with recombinant IFNγ (rIFNγ) to mimic elevated IFNγ signaling (Fig. 2A). rIFNγ addition at the start of stroma-supported organoid culture was sufficient to significantly disrupt young and old alveolar organoid formation (Fig. 2B-C), demonstrating repression of AT2 cell self-renewal by IFNγ independent of age. To investigate the impact of IFNγ on AT1 differentiation, we quantified organoids with RAGE+ AT1-like cells using immunofluorescent staining (Fig. 2D, Supp Fig. 2G). IFNγ-treated organoids showed a subtle trend of decreased RAGE+ AT1 cell differentiation with statistical significance in old but not young organoids. Alveolar organoids propagated from a conditional AT2-specific IFNγR1 knockout mouse did not have reduced organoid formation in response to rIFNγ (Fig. 2E-G) and were un-responsive to IFNγ-induced expression of the MHC-I component *B2m* (Fig. 2H), demonstrating that the effect of IFNγ on organoid growth operates through the IFNγR1 on AT2 cells.

A variety of lung airway progenitors are capable of alveolar repair (*8*), but it was not clear whether airway-derived progenitors are capable of alveolar repair in the IFNγ-rich environment of the aged lung. To model this, multi-potent bronchiolar progenitors were sorted (Supp Fig. 1A) from old and young mice for organoid culture and grown with rIFNγ (Supp Fig. 2H). The formation of surfactant protein C (SPC)+ alveolar organoids decreased significantly after IFNγ treatment in both old and young cells, whereas the number of bronchiolar organoids with SCGB1A1+ secretory cells was unaffected (Supp Fig. 2I-K). This parallels observations of AT2 cell, but not bronchiolar cell regenerative loss in aged lungs after injury (*3*).

To test whether IFNγ plays a causative role in aged AT2 cell regenerative decline *in vivo*, we utilized IFNγ neutralizing antibody treatment followed by organoid formation assays. Young and old mice were given anti-IFNγ neutralizing or IgG control antibodies for one week, before sorting of AT2 cells and subsequent organoid cultures (Fig. 2I, J). As shown previously, surface MHC-I was elevated in aged AT2 cells from IgG-treated old controls compared to young controls (Fig. 2K, L). AT2 cells from anti-IFNγ treated aged mice had significantly reduced surface MHC-I complex, indicative of dampened IFNγ signaling (Fig. 2K, L). AT2 cells from aged mice treated with anti-IFNγ had significantly improved organoid-forming efficiency compared to aged IgG controls (Fig. 2M), indicating that IFNγ signaling in aged lungs disrupts the regenerative capacity of AT2 cells reversibly. AT1 differentiation potential of organoids did not change (Fig. 2N), likely because AT1 differentiation begins after 7-10 days in stroma-supported organoids (*29*) and therefore potentially after an IFNγ-induced repressive cell state in aged cells has resolved *ex vivo*.

### T cell accumulation within tertiary lymphocyte structures is responsible for elevated IFNγ in the aged lung

To identify the cellular origin of IFNγ signaling in the aged lung, we utilized our scRNA-seq dataset of young and old whole murine lung aging focusing on the lymphocyte compartment (Fig. 3A, B, *26*) coupled with flow cytometry (Supp Fig. 3A, B). Old mouse lungs contained an elevated abundance of select T cell populations based on scRNA-seq milo analysis, including a unique population of GZMK+ T cells shown to be more frequent in the aged lung (Fig. 3C, *26, 36*). IFNγ transcripts were sparse but elevated in several aged T cell populations and found at the highest levels in aged natural killer (NK) cells, CD8+ cytotoxic T cells, and GZMK+ CD8+ T cells but absent in B and plasma cells (Fig. 3D). Congruently, we found that CD8+ T cells and NK cells from old human lung scRNA-seq data have an elevated IFNγ signature and that these CD8+ T cells have an increase in IFNγ expression in advanced age (Fig. 3E, F).

**Figure 3.**
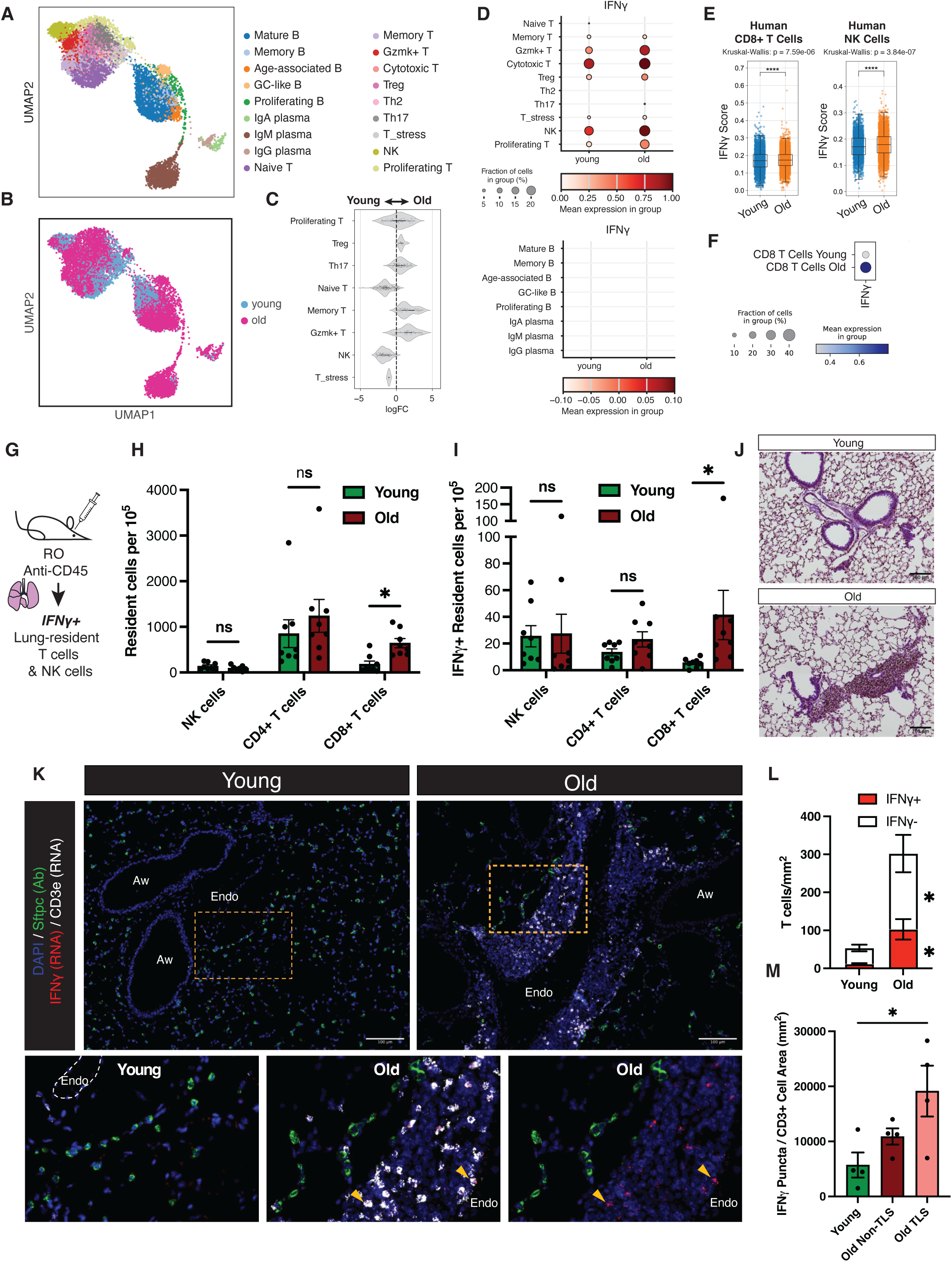
T cell accumulation within tertiary lymphoid structures is responsible for elevated IFNγ in the aged lung. A) UMAP representation of subclustered lymphocytes in aging mouse lung (18 months) and young controls (3 months) scRNA-sequencing without injury. Whole murine lung aging dataset originally made public in Gote-Schniering et al. (*26*). B) UMAP labeled by old or young lung samples and (C) T cell abundance fold change in aged mouse lung by annotated cell type using milo analysis, excluding cell types that did not meet the threshold for quantification. D) IFNγ transcript via dotplot in lung lymphocytes separated by age, grouped into plots T cell/NK cell or B cell populations, and broken down by annotated cell type. E) Human lung cell atlas (*23*) comparison of IFNγ response genes by GSEA of hallmark pathways in human CD8+ T cells and NK cells across age cohorts; young (<30 years), old (>60 years). F) A comparative dotplot of CD8+ T cell IFNγ gene expression in the same population of human lung cell atlas samples. G) Experimental diagram of retroorbital injection of anti-CD45 into old and young mice followed by lung flow cytometry to measure lung-resident lymphocyte populations for intracellular IFNγ. H) Abundance of key resident lung lymphocyte subpopulations and (I) resident IFNγ+ subpopulations. J) Gross pathology and immune cell infiltration shown by H&E stain in comparable old and young perivascular regions. K) Representative fluorescent imaging of bronchiolar/vascular regions in old and young lung using a combined immunofluorescence and RNAscope approach, including staining with anti-SPC for AT2 cells (green), IFNγ mRNA transcript (red), and CD3e mRNA transcript (white). Zoomed-in cutouts highlight IFNγ+ CD3e+ T cells within TLS in the distal lung, with examples marked by yellow arrows. “Aw” labels airways and “Endo” marks endothelial vessels. L) Quantification of total IFNγ+ CD3e+ T cells and CD3e+ T cells per mm^2^ in perivascular/peribronchiolar lung regions. M) Quantitation of the density of puncta marking IFNγ mRNA transcript in CD3e+ cells compared by age and localization within TLS-like structures. *P < 0.05, **P < 0.01, ***P < 0.001, ****P < 0.0001.

To further assess which lung-resident lymphocyte populations were responsible for elevated IFNγ release in aged lungs, we measured IFNγ by flow cytometry in old and young lung lymphocytes after excluding circulating lymphocytes that had been marked by a retroorbital injection of anti-CD45 antibody (Fig. 3G, Supp Fig. 3A, B). Lung-resident CD8+ T cells were more abundant in aged lungs compared to young lungs (Fig. 3H) and there was a significant increase in lung-resident IFNγ+ CD8+ T cells in the aged lung (Fig. 3I). To functionally test whether a lymphocyte population in the aged lung released more IFNγ when activated, we measured IFNγ release by flow cytometry after culture with Ionomycin and Phorbol 12-myristate 13-acetate (PMA) (Supp Fig. 3C). IFNγ+ resident T cells again made up a significantly higher proportion of lymphocytes isolated from aged lungs (Supp Fig. 3D, E), but no aging difference was observed in terms of lung-resident T cell activation as measured by IFNγ release (Supp Fig. 3F). Aged lung NK cells tended to be more frequently IFNγ+ than young NK cells after stimulation without reaching significance (Supp Fig. 3F).

We next sought to localize IFNγ+ T cells in the aged lung to understand how they could impart an effect on AT2 cells. An assessment of healthy aged lung pathology revealed striking perivascular and peribronchiolar cellular aggregates with severity that varied across animals (Fig. 3J). Recent reports in the healthy aged lung have identified these as tertiary lymphoid structures (TLS, *26*, *37, 38*) made up of a defined mixture of specialized endothelial, myeloid, and lymphoid cells, including GZMK+ CD8+ T cells. We hypothesized that T cells releasing IFNγ may accumulate in these structures in old lungs. Using RNA in situ hybridization we observed elevated IFNγ transcript in TLS areas in old mice and not in similar peribronchiolar/perivascular regions in young mice (Fig. 3K, Supp Fig. 3G, H). IFNγ transcripts in aged lung were almost exclusively co-localized with *CD3e* transcript labeling T cells, which accumulate in TLS, and IFNγ+ T cells were found in elevated abundance in aged perivascular regions (Fig. 3L, Supp Fig. 3I). Aged T cells within TLS regions expressed significantly elevated levels of IFNγ transcript compared to young T cells (Fig. 3M). *Ncr1*+ NK cells were also detected in old and young lungs but were not specifically localized to TLS (Supp Fig. 3G), had similarly low numbers in both old and young lungs (Supp Fig. 3J, K), and had no statistically significant difference in IFNγ transcript density in the aged lung (Supp Fig. 3L). Though we saw limited evidence of direct T cell interactions with AT2 cells in distal alveolar regions, SPC+ cells appeared densely concentrated at the TLS boundary in aged lungs (Fig. 3K, Supp Fig. 3H). These observations raise the possibility of a spatial gradient with a functional distinction between AT2 cells closely interacting with IFNγ+ T cells near TLS and more distally located AT2 cells further from IFNγ release.

We asked whether IFNγ+ T cells were required for the elevated IFNγ response in aged AT2 cells using young and old *Foxn1* KO mice which lack T cells. Aged *Foxn1* KO mice lacked elevated MHC-I complex on the surface of AT2 cells (Supp Fig. 3M), reflective of a low IFNγ signaling environment. Alveolar organoids grown from old and young *Foxn1* KO mice showed no significant difference in organoid-forming efficiency, in contrast to AT2 cells from aged C57BL/6 mice (Supp Fig. 3N, Fig. 2A-C).

### Immunoproteasome function is required for the IFNγ-rich microenvironment in the aged lung and contributes to aged AT2 regenerative decline

We next aimed to explore the impact of reduced MHC-I antigen presentation in the aged lung microenvironment on aged AT2 cell regenerative properties. Using mice with a triple knockout of all three catalytic subunits of the immunoproteasome, i.e. *Psmb8*, *Psmb9*, and *Psmb10* (IP KO) provided a means, as IP KO mice have impaired MHC-I antigen presentation to CD8+ T cells and therefore compromised CD8+ T cell development (*39*). AT2 cells sorted from aged IP KO mice had partially rescued alveolar organoid formation and diameter compared to wildtype aged mice in feeder-free organoid assays (Fig. 4A-C), indicating a crucial role for immunoproteasome function and/or antigen presentation in aged AT2 regenerative decline. Aged IP KO AT2 cells showed reduced expression of the MHC-I complex component *B2m* by RTqPCR (Fig. 4D), confirming disruption in age-related chronic interferon signaling. Furthermore, aged IP KO alveolar organoids had an elevated ratio of Ki67+ proliferating cells compared to age-matched wildtype alveolar organoids (Fig. 4E, F). AT2 cell density in aged IP KO mice trended upwards compared to aged controls (Supp Fig. 4A-B).

**Figure 4.**
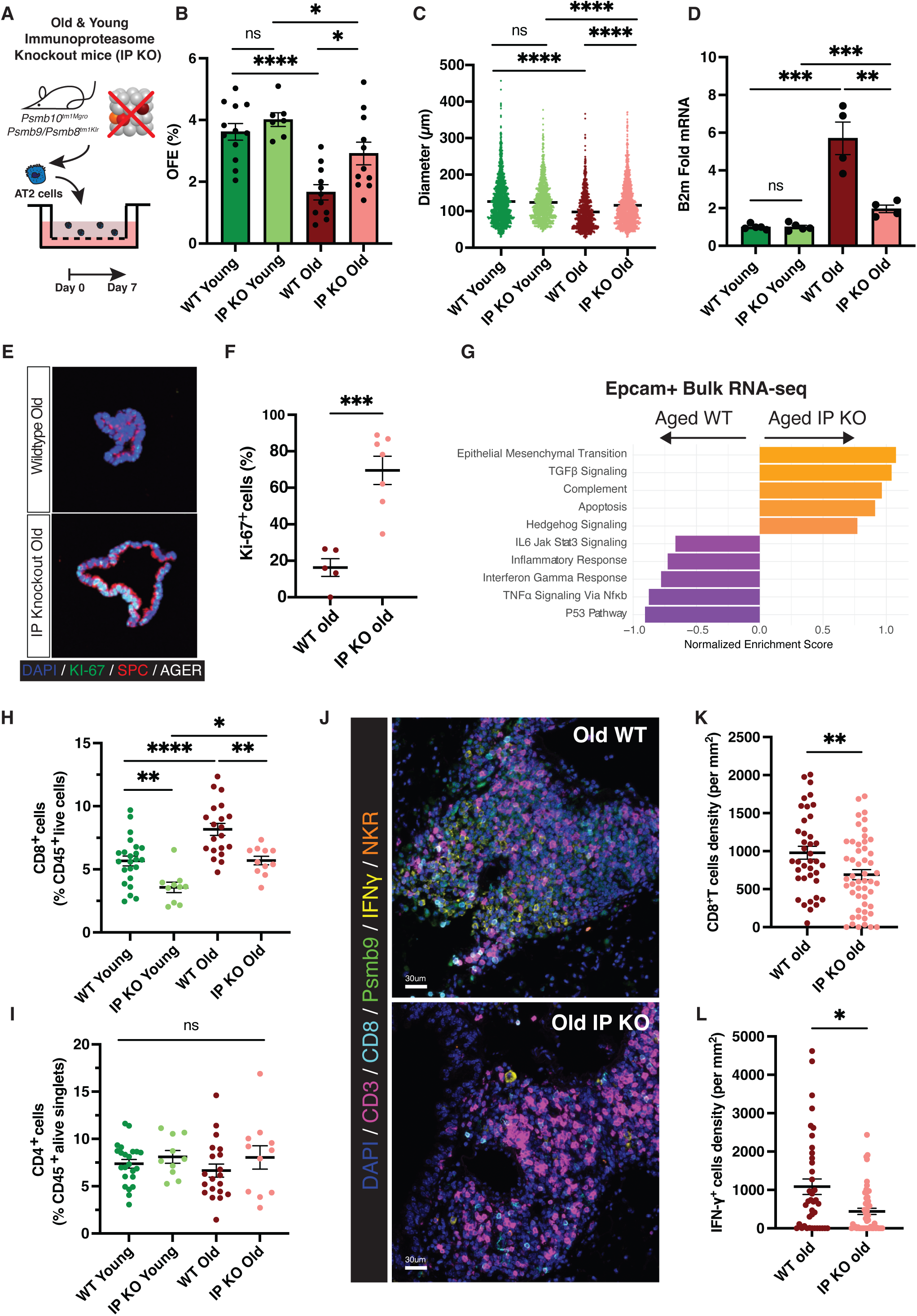
**Immunoproteasome is required for aged AT2 cell regenerative decline and the establishment of IFN**γ+ **T cells in tertiary lymphoid structures.** A) Schematic figure to show the culture of alveolar organoids from young or aged immunoproteasome knockout (IP KO) and control C57BL/6 (WT) mice in feeder-free media at 7 days of growth, including B) organoid forming efficiency (OFE) and (C) diameter. D) RTqPCR of the IFNγ-inducible MHC-I complex component B2m in isolated lung AT2 cells from young or aged immunoproteasome KO and WT mice. E) Representative immunofluorescence staining of Ki-67 to mark proliferating cells with AT2 cell marker SPC and AT1 cell marker Rage/Ager in corresponding feeder-free organoids from aged mice in A-C. F) The ratio of Ki-67+ proliferating cells in aged immunoproteasome knockout (IP KO) versus WT mice. G) Dysregulated pathways in aged immunoproteasome KO epithelium in comparison to WT based on DEG genes (padj < 0.05, LogFC <0), with bar size representing enrichment score. H) Flow cytometry quantifications of CD8+ and CD4+ (I) T cells as % of all CD45+ immune cells in aged immunoproteasome KO and WT control lungs. J) Zoomed-in multiplex fluorescent images of TLS in aged lungs of immunoproteasome KO and WT controls with K) corresponding absolute quantification of CD8+ T cell and IFNγ+ cell (L) densities. *P < 0.05, **P < 0.01, ***P < 0.001, ****P < 0.0001.

We considered whether loss of immunoproteasome function and CD8+ T cell interactions disrupt the formation of an IFNγ-rich lung microenvironment. Bulk RNA sequencing of EPCAM+ lung epithelial cells from aged IP KO mice, containing mostly AT2 cells, confirmed a reduced IFNγ response, whereas TGFb and hedgehog signaling pathway components were increased (Fig 4G, Supp Fig. 4C-D). Flow cytometry of digested lungs revealed reduced proportions of NK and CD8+ T cells in lung tissue, spleen and lymph nodes of aged IP KO mice compared to aged controls, whereas other lymphoid and myeloid populations remained stable (Fig. 4H, I, Supp Fig. 4G-I). Aged IP KO lungs retained TLSs (Supp Fig. 4E), so we assessed their cellular composition and IFNγ release. In accordance with our flow cytometry data, aged IP KO lungs showed significantly reduced density of CD8+ T cells and IFNγ+ cells within TLS by multiplex immunofluorescence staining (Fig. 4J-L). Taken together, our data indicate that the formation of an IFNγ-rich aged lung microenvironment is dependent on immunoproteasome function during the aging process and furthermore point to a role for a feedforward cycle of elevated IFNγ, enhanced antigen presentation, increased T cell accumulation, and therefore further elevated IFNγ signaling contributing to the decline in AT2 cell functions during aging.

## Discussion

Our study establishes that local IFN*γ* signaling in homeostatic aged lungs actively alters AT2 cell state and represses alveolar regeneration. Chronic inflammatory signaling is a hallmark of aging commonly thought to be a byproduct, not a cause, of age-related decline (*10,41,42*), yet our findings are consistent with an immunological theory of aging (*43*). We show that aged lungs have a chronic IFN*γ*-rich inflammatory environment with elevated abundance of resident CD8+ T cells, independent of infection or injury. Neutralization of IFN*γ* signaling or disrupting the formation of an IFN*γ*-rich microenvironment via deletion of the immunoproteasome partially rescued the organoid-forming efficiency of aged AT2 cells, highlighting the impact of epithelial-immune communication during lung aging. Our results demonstrate that aged AT2 cells are more sensitive to subtle inflammatory processes in the local microenvironment than previously thought (*10*), and that improved aged AT2 cell-directed regeneration is plausible despite the accumulation of mutations and transcriptional changes seen in aged AT2 cells (*41, 44*).

In agreement with our findings in aged samples, interferon signaling in the lung has previously been shown to have a broad impact on lung architecture and effects on AT2 cell state. Direct instillation of IFNγ (*45*) or elevated IFNγ from lung tissue-resident memory (TRM) lymphocytes (*46*) contributes to loss of alveolar density in young mice. This mirrors the reduced alveolar density seen during aging (*47,48*) and the more severe emphysema in COPD (*49,50*). We here show that aged AT2 cells shift towards an MHC-I-high state, which shares similarities with cell states identified after injury or infection, namely the activated or primed AT2 state (*11, 27*) and an ISG-high state driven by Type I interferon after influenza challenge (*16*). Aged AT2 cells are, however, transcriptionally distinct. In particular, the chronic IFNγ response we observe in aged AT2 cells lacks expression of some ISGs, such as *Irf1*, which may be the result of known negative feedback mechanisms due to the chronic nature of the IFNγ exposure. For example, we observed elevated *Socs2* in aged AT2 cells, a negative feedback inhibitor of the JAK/STAT pathway (Fig. 1J, *35*). The longtime accepted paradigm that IFN signaling is protective against infection and cancer at the cost of cell growth (*18*), is reflected in our findings, though this has been challenged as context dependent. Some human alveolar organoid culture conditions appear to overcome the repressive effects of added rIFNγ (*15*) and in a mouse influenza model AT2 cells recover from an IFN-responsive repressed state through Oncostatin-M signaling (*16*). These studies raise questions about whether some mechanisms that dictate inflammatory resolution have been lost in aged cells.

Our work assesses why aged AT2 cells are less poised for self-renewal in the healthy aging lung, aligning with recent studies that have dissected poor alveolar injury resolution in aged lungs (*26,31*). We observe the formation of an inflammatory niche due to IFN*γ*+ CD8+ T cells accumulation in TLS, which a recent study showed is driven by a fibroblast-macrophage-T cell axis during aging (*54*). In aged mice, T cell-derived GZMK and inflammatory macrophage-derived IL-1β are thought to play a role in the persistence of the *Krt8*^hi^ AT2-AT1 transitional state after bleomycin (*26*) or influenza challenge (*31*) respectively. We found that loss of CD8+ T cells and the corresponding loss of IFN*γ* release in aged mice lacking *Foxn1* or the immunoproteasome rescued AT2 cell organoid formation. Our results in combination with this work suggest that age-related CD8+ lung-resident T cells can disrupt alveolar regeneration with several independent but interrelated mechanisms (IFNγ, GZMK, and IL-1β), highlighting the complex impact of age-related inflammation on AT2 cells.

Our data suggest that aged AT2 regenerative decline is partially reversible and dependent on immunoproteasome function. Ongoing clinical testing of specific immunoproteasome inhibitors with minor side effects may prove useful for treatment of age-related lung disease. Restricting IFNγ release from T cells or depletion of IFNγ-releasing T cells in the aged lung serves as a promising candidate to re-establish alveolar repair in age-related chronic lung disease, though this will need to be balanced with their protective role in the immunosurveillance of lung malignancies and infections associated with aging (*51*). Our findings reveal an opportunity for more targeted therapeutic approaches to intervene in susceptibility to age-related lung disease in the near future.

## Acknowledgements

We thank Ron Mathieu, Mahnaz Paktinat, Tyriq Jones, Betelhem Gemechu, Ranjan Maskey, and others at the BCH flow cytometry core for their technical support and phenomenal dedication to our experiments. We thank past and present members of the Kim, Meiners, and Lehmann laboratories, Judith Agudo and Marcia Haigis for important discussions and varying forms of support as this work developed. We thank Frauke Koops, Gesine Rode, Ulrike Büchsler and Jasmin Tiebach, the histology and flow cytometry units of the Research Center Borstel for excellent technical support. We acknowledge the support of Ulrike Seifert and Clemens Cammann from the University Greifswald and Regeneron for supplying IP KO mice (Triple Immunoproteasome KO Mice (*Psmb8, Psmb9, Psmb10* KO) - VG1230/VG11677). We acknowledge COPD-iNET (www.copd-inet.com) for fostering scientific exchange and collaboration on this topic.

## Author contributions

J.J. designed, conceived, performed, and analyzed experiments, and wrote the manuscript. K.G. and J.G.S., designed, conceived, performed, and analyzed experiments, and co-wrote the manuscript. Z.O., J.W., M.P., M.S. designed and performed laboratory experiments and analyzed data. J.G., M.C.M., G.B., A.C., J.J., and P.P. generated RNA/ATAC sequencing data in the laboratory. J.G.S., M.M., S.H.S., M.C.M., Q.H., J.J., and P.P. analyzed sequencing data. M.L., S.M., and C.F.K. supervised the design of the study, conceived experiments, acquired funding, and co-wrote the manuscript. All authors reviewed and edited the manuscript.

## Data and materials availability

## Supplementary Materials

Materials and Methods

Figs. S1 to S4

References (*51–56*)

Data S1

## Materials and Methods

### Mice

Aged C57BL/6J mice (18-24 months) were procured from Charles River Laboratories or Jackson laboratory then aged further within the Boston Children’s Hospital mouse facility, while young C57BL/6J (2-4 months) were purchased from a matching supplier for experiments. IFNγR1 Cre-conditional knockout mice (Jackson Laboratory #025394) on a C57BL/6J background were crossed to Sftpc-CreER (Jackson Laboratory #025394) and R26-LSL-TdTomato (Jackson Laboratory #007914) to create TdTomato-labeled AT2 cells without functional IFNγR. All mice were maintained and bred in under specific pathogen-free (SPF) conditions and housed in groups. All experiments were approved by the Boston Children’s Hospital Animal Care and Use Committee. Roughly equivalent numbers of age-matched male and female mice were used in experiments where possible, with no clear sex differences observed in the experiments conducted. For immunoproteasome KO experiments, Wild-type C57BL/6J mice and B6.129-Psmb10^tm1Mgro^ Psmb9/Psmb8^tm1Klr^ (*46*) on a C57BL/6J background were used in accordance with institutional and governmental regulations. Mice were bred and aged under specific pathogen-free (SPF) conditions at the animal facility of the Research Center Borstel. During experiments, animals were housed in type II long IVCs with wood bedding (LTE E-001, Abedd, Latvia) at 20-24°C, 45-65% relative humidity, and a 12h light/dark cycle. γ-Irradiated maintenance diet (ssniff Spezialdiäten GmbH, Germany) and water were provided *ad libitum*. Each cage contained nesting material and red plastic boxes for environmental enrichment.

### Anti-IFNγ neutralizing antibody treatment

To ablation of IFNγ signaling in the lung micronenvironment, anti-IFNγ antibodies (BioXCell #BE0055) and IgG isotype control (BioXCell #BE0088) were diluted to 1μg/1μl in sterile PBS prior to intraperitoneal injection (IP) and intratracheal instillation (IT). Mice were administered either anti-mouse IFNy or IgG control by IP (100 μg) and by IT (50 μg). Antibodies were administered three times, once every third day, for a total of six days. One day after the final treatment, mice were euthanized and their lungs processed for FACS, or one week after the first treatment.

### Flow cytometry and fluorescence-activated cell sorting of lung epithelium

To isolate AT2 cells and measure surface MHC-I complex, murine lungs were harvested for FACS as previously described (*3,63*) with minor modifications. Briefly, mice were euthanized by avertin overdose and lungs were flushed with cold PBS via cardiac perfusion. Lungs were inflated with dispase (Corning, USA) with an intratracheal syringe, then lungs were excised, transferred to ice, and minced. This was followed by enzymatic digestion of lung tissue in PBS containing 0.0025% DNase (Sigma-Aldrich, USA) and 100 mg/mL collagenase/dispase (Roche, Switzerland) for 45 min at 37°C with gentle agitation on a rotator. The resulting suspension was passed through 100- and 40-μm cell strainers (Falcon, USA) and pelleted by centrifugation at 1000 rpm for 5 min at 4°C. The pellet was resuspended in red cell lysis buffer (0.15 M NH4Cl, 10 mM KHCO3, 0.1 mM EDTA) for 90 s at room temperature before quenching with DMEM (Gibco) containing FBS. Following a second centrifugation step at 1000 rpm for 5 min at 4°C, cells were resuspended in PBS with 10% FBS for immunostaining. The antibody panel included anti-CD31 APC (Thermo Fisher Scientific #BDB559864), anti-CD45 APC (Thermo Fisher Scientific #BDB551262), anti-CD326 (EpCAM) PE/Cy7 (Biolegend #118216), anti-Ly-6a/e (Sca1) APC/Cy7 (Thermo Fisher Scientific #560654), anti-CD24 (BD Biosciences #553261), and anti-H2-K^b^ (Thermo Fisher Scientific #11-5958-82) all at 1:100. Dead cells were excluded using DAPI (Sigma-Aldrich) and all antibodies had single stain and FMO controls in each experiment. AT2 cell (CD31-CD45-EpCAM+Sca1-CD24-) and bronchiolar progenitor (CD31-CD45-EpCAM+Sca1+CD24low) sorting was carried out on a FACS Aria II (BD Biosciences) and data analysis of H2-K_b_ levels was performed in FlowJo v10.

### Flow cytometry of lymphocytes

For immunoproteasome KO immune cell characterization, mice were euthanized by CO₂ inhalation. Following disinfection with 70% ethanol, skin was cut, blood was collected via cardiac or portal vein puncture and immediately mixed with 2.7% EDTA in PBS if immune cell analysis was strived. The pleural cavity was opened, and the lungs were perfused through the heart with 20 mL pre-warmed PBS to minimize contamination of lung-resident immune cells by blood-derived immune cells. The thymus and lung-draining lymph nodes were carefully separated from the lung tissue. Axillary, brachial, and mesenteric lymph nodes and spleen were dissected. All organs were kept on ice during processing. Lymph nodes and spleens were mechanically dissociated by gently grating through a 100 µm mesh, followed by washing with MACS buffer (5% fetal bovine serum in PBS). Perfused lungs were minced and digested in 10 mL enzyme mix containing 30 µg/mL deoxyribonuclease I (D4527, Merck, Germany) and 0.7 mg/mL collagenase (C5138, Merck, Germany) in RPMI 1640 supplemented with 10% FBS, 1 mM sodium pyruvate, 100 U/mL penicillin, 0.1 mg/ mL streptomycin, 10 mM HEPES, and 50 µM 2-mercaptoethanol. Digestion was performed for 30 min at 37 °C with agitation (180 rpm). Cell suspensions were filtered through 100 µm strainers (Corning) and washed by centrifugation at 427 × g for 5 min at 4 °C in HBSS containing 3% FBS. Erythrocytes were lysed using Ery-Lysis buffer (0.15 M NH₄Cl, 10 mM KHCO₃, 0.1 mM EDTA, pH 7.4). After lysis, cells were washed with 30 mL and resuspended in 3 mL of 3% FBS in HBSS. The suspension was overlaid on 3 mL Lympholyte separation medium (Cedarlane, Canada) and centrifuged at 1.200 × g for 20 min at room temperature (without brake). Mononuclear cells were collected from the interphase and washed with 5 mL 3% FBS in HBSS. Cells were resuspended in FACS buffer (2% FBS, 0.1% NaN_3_, 0.2mM EDTA in PBS containing Fc-block (BioLegend)) for staining and flow cytometric analysis. Final cell suspensions were passed through 30 µm strainers (Sysmex Partec GmbH, Germany) to remove debris. Finally, stained cells were measured by a flow cytometry (LSR II, BD Bioscience, USA) and analyzed in FlowLogic v8. LIVE/DEAD™ Fixable Blue Dead Cell Stain Kit (L23105, Thermo Fisher, USA) was used to determine the cell viability. Antibodies used in flow are listed below:

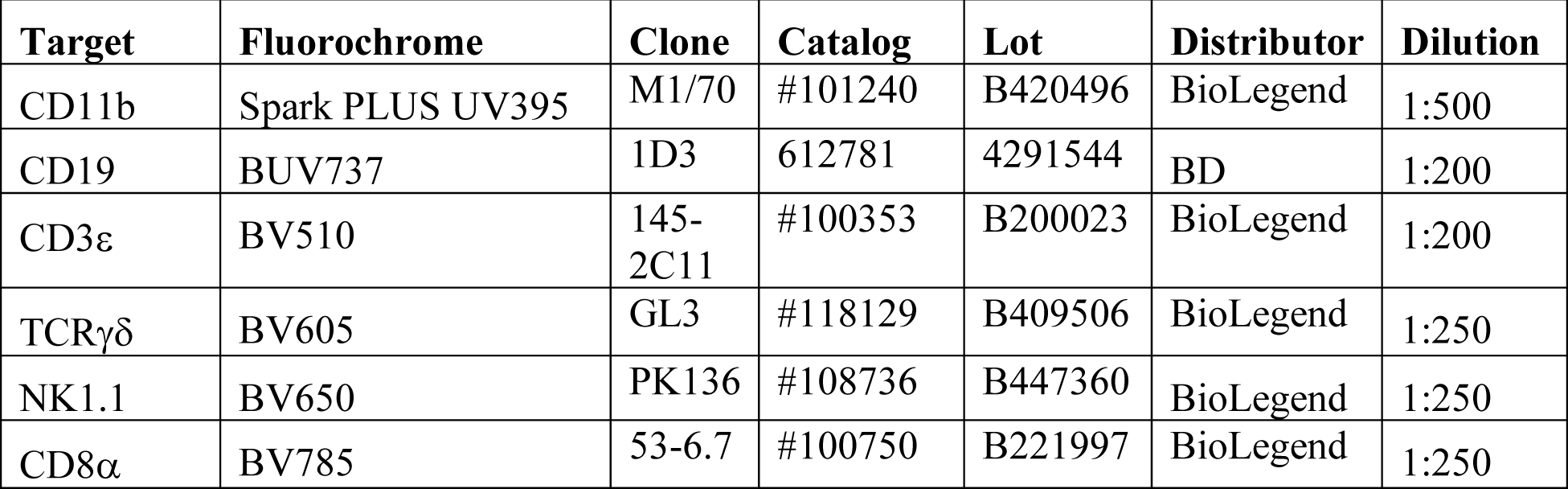
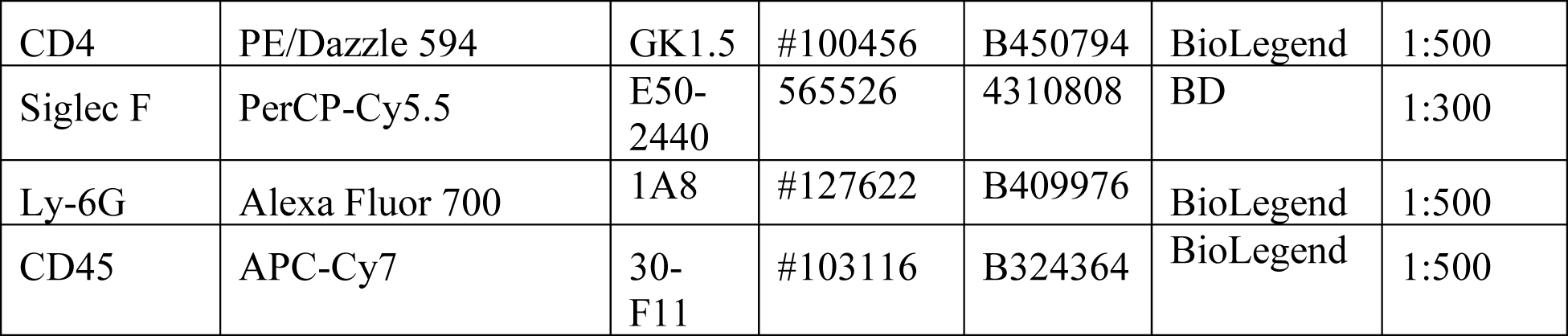

### Flow cytometry of intracellular IFNγ

To measure intracellular IFNγ in mouse lung-resident lymphocytes, mice received a retro-orbital injection of anti-CD45 BV750 (1:10 in sterile PBS, 100 µL per mouse) to label circulating immune cells immediately following avertin anesthesia. One mouse was not injected as an unlabeled control. Bronchoalveolar lavage (BAL) was performed using two sequential passes of 1 mL and 500 µL of cold BAL buffer (PBS supplemented with 2% FBS and 0.4 mM EDTA) and saved on ice. Lungs were then processed as described above, with the following modifications: lung inflation and digestion were performed using HBSS containing 100 µg/mL Liberase and 0.025 mg/mL DNase. Following red blood cell lysis, cells were either fixed and immunostained right away or live lymphocytes were enriched in a 40/90 Percoll density gradient (*64*) for *ex vivo* culture. Briefly, cells were resuspended in cold DMEM with 40% Percoll and underlaid with 90% Percoll to achieve a 40/90% discontinuous gradient, then centrifuged at 2,500 rpm for 20 min with slow acceleration and deceleration. Lymphocytes were harvested from the interface, washed in HBSS, and resuspended in T-cell medium (RPMI-1640 supplemented with 10% FBS, 20 mM HEPES, 1 mM sodium pyruvate, 2 mM L-glutamine, 50 µM β-mercaptoethanol, 1× non-essential amino acids, and 1× penicillin/streptomycin). Cells were incubated for 3 h at 37°C in either control medium, unstimulated media with the Golgi export inhibitors Brefeldin A (BD Biosciences #555029) and 2 µM Monensin (Biolegend #420701) or stimulated medium with Cytek Cell Stimulation Cocktail [PMA, Ionomycin, Brefeldin A, and Monensin] (Cytek Biosciences TNB-4975). Cells were stained with LIVE/DEAD Fixable Violet (Thermo Fisher Scientific L34955) to exclude dead cells, then fixed and permeabilized using the BD Cytofix/Cytoperm kit (#554714) or Foxp3/Transcription Factor Staining Buffer Set (Thermo Fisher Scientific #00-5523-00) per the manufacturer’s instructions. After an initial incubation with Fc block (Biolegend #101302) in flow staining buffer on ice, intracellular and surface staining was performed overnight at 4°C with anti-CD45 FITC (BD Biosciences #553080), anti-CD3 APC-Cy7 (Biolegend # 100222), anti-CD4 PE-Cy7 (Biolegend #100528), anti-CD8 APC (Biolegend #100712), anti-NK1.1 PE (Biolegend #156504), and anti-IFNγ BV605 (Thermo Fisher Scientific #406-7311-82) all at a dilution of 1:100. Single stain and FMO controls were prepared for each antibody. Cells were washed and resuspended in PBS with 10% serum for flow cytometry acquisition on a BD LSRFortessa and analysis in FlowJo v10 (BD).

### Magnetic bead sorting isolation of lung epithelium

Murine distal lung epithelial cells (CD45−/CD31−/EpCAM+) were obtained after enzymatic and mechanical digestion of mouse lungs as previously described (*6*). Briefly, mice were euthanized by overdose of ketamine (150 mg/kg) and xylazine (10 mg/kg) and followed by lethal blood withdrawal at the inferior vena cava. Blood was flushed from the lungs with PBS, and the lungs were inflated intratracheally with Dispase (Corning) and inflated with 1 % low gelling temperature agarose (Sigma-Aldrich). The lungs were excised and incubated in Dispase (Corning) for 40 mins at room temperature. Lungs were then minced and filtered through 100 µm and 40 µm cell strainers (Falcon). Using magnetic bead sorting (MACS), macrophages and white blood cells as well as endothelial cells were removed using CD45 and CD31 beads (Miltenyi Biotec), respectively. Epithelial cells were enriched using CD326 (EpCAM) beads (Miltenyi Biotec) according to the manufacturer’s instructions. The enriched EpCAM+ cells were snap-frozen in liquid nitrogen and stored at −80°C until RNA isolation for bulk RNA sequencing. To magnetically sort AT2 cells for organoid cultures in immunoproteasome knockout mouse experiments (Figure 4), a different protocol was used. After euthanasia, a volume of 1.5 ml Dispase (354235, Corning, USA) was intratracheally injected to inflate the mouse lung. Subsequently, lung lobes were removed and transferred to a sterile cell culture dish for further digestion at RT for 45 min. Next, lungs were minced into small pieces using a scalpel and digested with lung lysis buffer (1× Antibiotic-Antimycotic (15240062, Gibco, USA) buffered DMEM/F12 medium supplied with 450 U/ml collagenase (17100017, Thermo Scientific, USA), 5 U/ml Dispase, 50 U/ml DNase (D5025, Merck, Germany)) at 37°C for 35 min. The cell suspension then was neutralized by adding an equal volume of MACS buffer (5% FBS in PBS) and filtered through the 70 μm cell strainer to remove undigested tissues. After lysis of erythrocytes using ACK buffer (0.15 M NH_4_Cl, 10 mM KHCO_3_ and 0.5 M EDTA), cells were resuspended with 10% FBS supplied DMEM/F12 medium and added to a plate coated with purified anti-mouse CD31 (102502, BioLegend, USA)/purified anti-mouse CD45 (157602, BioLegend, USA) antibodies for 1 h at 37°C to exclude endothelial and immune cells. From the CD31^-^/CD45^-^ lung cell fraction, Epcam-positive cells were labeled with an anti-mouse Epcam antibody directly conjugated to magnetic beads (Miltenyi Biotech #130-105-958) and purified by using the AutoMACS Pro Separator (Miltenyi Biotech #130-092-545) with positive selection program. To isolate epithelial cells from alveolar organoids embedded in Matrigel, a digestion of 2 mL of TrypLE (Thermo Fisher Scientific #12604039) per 100 μL Matrigel plug was used. The preparation of these cultures is described below. After 15 minutes at 37 °C with intermittent pipetting to mechanically disrupt the organoids, it was confirmed that organoids had become a single-cell suspension using a hemocytometer before quenching with 10% FBS in PBS. Organoid epithelial cells were then enriched using CD326 (EpCAM) beads (Miltenyi Biotech #130-105-958) according to the manufacturer’s instructions to remove stromal cells before RNA isolation. Feeder-free organoids in Supp Fig 2 were isolated in the same manner using TrypLE digestion without a magnetic cell isolation step.

### RNA isolation from sorted cells

AT2 cells isolated from mouse lung or organoids, between ∼10,000 and ∼100,000 cells were pelleted before the addition of lysis buffer from the absolutely RNA microprep kit (400805, Agilent technologies, USA) and freezing at −20 °C. After thawing, RNA was isolated following manufacturer’s instruction, eluted in 30 µl elution buffer, and storage at −80 °C. For RTqPCR of wildtype and immunoproteasome knockout mice (Figure 4), RNAs of isolated AT2 cells were extracted by using NucleoSpin RNA Plus XS kit (740990.50, Macherey-Nagel, Germany) following manufacturer’s instruction. For bulk RNA sequencing, AT2 cells from young (7 weeks) and old (18 months) mice were isolated by FACS of CD31-CD45-Epcam+Sca1− cells as described above. RNA was isolated using an Absolutely RNA Kit (Agilent technologies, #400800) according to the manufacturer’s instructions. For bulk RNA sequencing of epithelial cells from aged wildtype versus aged immunoproteasome knockout mice, cryostored magnetically sorted EpCAM^+^ cells were lysed using RLT Plus Lysis Buffer (Qiagen). RNA isolation was performed using the RNeasy Mini Kit (Qiagen) according to the manufacturer’s instructions and eluted in 40 µl water. RNA concentration was determined using a Nanodrop 1000 and samples were stored at −80 °C until sequencing.

### Real-time quantitative PCR

For RTqPCR of AT2 cells isolated by FACS from mouse lung or most organoid experiments, template RNA was used to synthesize cDNA with the High-Capacity cDNA Reverse Transcription Kit (Thermo Fisher Scientific #4374966). Relative expression of genes in AT2 cells was determined by using Taqman-based real-time quantitative PCR (RTqPCR) in a StepOnePlus real time PCR system. cDNA was used for pre-validated Taqman gene expression assays from Thermo Fisher Scientific include *Nlrc5* (Mm01243039_m1), *B2m* (Mm00437762_m1), *H2-k1* (Mm01612247_mH), *Psmb8* (Mm00440207_m1), *Psmb9* (Mm00479004_m1), and *Nrn1* (Mm00467844_m1). Relative expression of target genes was measured and expressed as 2^-ΔΔCt^ after normalizing to the housekeeping gene GapDH. For young or aged AT2 cells from wildtype versus immunoproteasome experiments specifically (Figure 4), SYBR Green based real-time quantitative PCR (qPCR) was used in a LightCycler 480 (Roche, Switzerland). To prepare the cDNA, 100 ng RNA was used as a template to synthesize cDNA using the Maxima First Strand cDNA Synthesis Kit (K1642, Thermo Scientific, USA). In brief, RNA was diluted in nuclease-free water and filled up to 14 μL, then the diluted RNA was mixed with 2 μL Maxima Enzyme Mix and 4 μL 5× Reaction Mix. The RNA mixture was subjected to incubation in a thermocycler (5331, Eppendorf, Germany) with 25°C for 10 min, 50°C for 15 min and 85°C for 5 min for cDNA synthesis. In short, synthesized cDNA was incubated with specific primers and SYBR Green Mix (04887352001, Roche, Switzerland) and subjected to repeated denaturation and annealing cycles. Relative expression of target genes was measured and expressed as 2^-ΔΔCt^ after normalizing to the housekeeping gene RPL19. Primer sequences used in this study are the following:

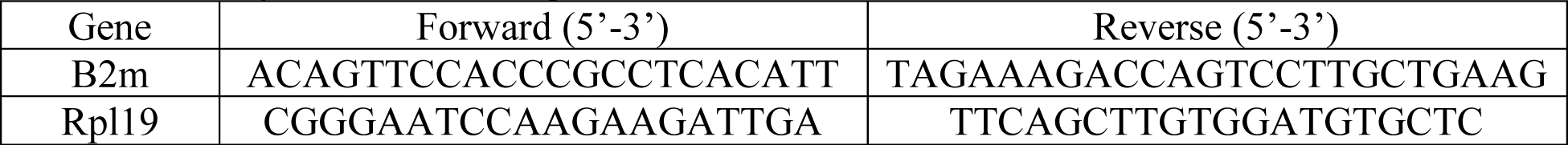

### Bulk RNA sequencing and analyses

For bulk RNA sequencing of young (7 weeks) and old (18 months) AT2 cells, isolated RNA were sent to the Molecular Biology Core Facilities at Dana-Farber Cancer Institute for rRNA depletion, quality control, library preparation, sequencing, and analysis. rRNA depletion was done using the Qiagen FastSelect rRNA depletion kit, libraries were prepared with Roche Kapa Biosystem RNA Hyperprep reagents, and those libraries were quality controlled via Qubit and Tapestation 4200. Paired-end reads (150 bp) were sequenced on an Illumina NovaSeq X Plus platform, aligned to the mm10 reference genome using STAR with gene annotations from Ensembl. Post-alignment quality control was performed using RSeQC, including assessment of read distribution across genomic features, ribosomal RNA (rRNA) depletion efficiency, and gene body coverage uniformity. Sample identity was confirmed by pairwise correlation of SNP profiles derived at HLA loci using snp. Transcript- and gene-level expression were quantified using Cufflinks. Sample relationships were explored by principal component analysis and hierarchical clustering of Spearman rank correlations. Differential gene expression analysis was performed using DESeq2, where genes were considered differentially expressed at an adjusted p-value of p < 0.05 and a minimum absolute log₂ fold change of 0.3. Overlapping aged AT2 cell markers were assessed in comparable aged AT2 cell single cell sequencing datasets, including McQuattie-Pimentel et al. 2021 (FDR *q* < 0.05) and Angelidis et al. 2019 (FDR *q* < 0.1), then compared by Venn diagram. For bulk RNA sequencing of epithelial cells from aged wildtype versus aged immunoproteasome knockout mice, samples were sent to Novogene for quality control, library preparation, and sequencing. Messenger RNA was purified from total RNA using poly-T oligo-attached magnetic beads. After fragmentation, the first strand cDNA was synthesized using random hexamer primers followed by the second strand cDNA synthesis. The library was ready after end repair, A-tailing, adapter ligation, size selection, amplification, and purification. The library was checked with Qubit and real-time PCR for quantification and bioanalyzer for size distribution detection. Non-stranded libraries were prepared with the Novogene NGS RNA Library Prep Set (PT042) and sequenced at 6G/sample on the NovaSeq Xplus platform (25B flow cell) with PE150, 300 cycles (Read1: 150, Index i7: 10, Index i5: 10, Read2: 150). Raw reads were trimmed, filtered and mapped using the CLC Workbech software v. 24 (Qiagen) with default settings and inclusion of broken pairs. Reads were mapped against GRCm39 (mouse). Raw counts were corrected for batch bias using ComBat_seq (sva v.3.50.0) in R version 4.3.1. Then, differential expression analysis was done using the DESeq2 package (pAdjustMethod = “BH”, alpha = 0.05, v.1.42.1) and shrinkage of the Log-fold change (LFC, apeglm method) (Zhu, et al, 2018). Differentially expressed genes (adjusted p-value< 0.05, LFC>0) were extracted and used for downstream analysis and data exploration. Gene set enrichment analyses were performed using fsgea (1.28.0) package based on the Hallmark mouse gene set collection from MSigDB v2023 (mh.all.v2023) (*60*).

### Human lung single-cell RNA sequencing analysis

Gene signature scores were calculated using the Human Lung Cell Atlas (HLCA, *23*). To minimize potential confounding effects, cells derived from diseased individuals or smokers were excluded from the analysis. Donors were stratified by age, with individuals ≤35 years defined as young and individuals ≥60 years defined as old. Gene signature scores were computed at the single-cell level using the scanpy.tl.score_genes function implemented in Scanpy (version 1.11.4). This method calculates the average expression of genes within a predefined gene set and subtracts the average expression of a set of reference genes with matched expression distributions. Three biological pathways were evaluated in this analysis, including interferon-γ response (IFNγ), MHC class I antigen presentation, and immunoproteasome activity. Gene sets used to calculate these scores were derived from the following curated gene lists and their human homologs: IFNγ response genes (Data S1), MHC class I genes (*65*), and immunoproteasome gene list (Data S1). Gene scores were computed for each cell and subsequently compared between age groups and across CD4 T cells, CD8 T cells, NK cells, and AT2 cells.

### Re-analysis of murine lung aging single-cell RNA sequencing

For re-analysis, the original dataset from Gote-Schniering et al. (*26*) was subsetted to include only uninjured lung samples from day 0. No repeated preprocessing or additional filtering was applied. All subsequent analyses were conducted based on the cell type annotation provided in the original publication. An MHC-I complex score was calculated from the expression of B2m, H2-D1, and H2-K1 using scanpy’s sc.tl.score_genes() function with default settings (scanpy v1.12) for AT2 cells. Based on the resulting score distribution, cells within the top 25% were assigned to the high group (threshold = 0.1), whereas all remaining cells were assigned to the low group. Differential expression analysis was carried out with scanpy’s rank_genes_groups function using the Wilcoxon rank-sum test. Genes with an absolute log2 fold change greater than 0.5 and an adjusted *p* value < 0.05 were considered significant. Pathway enrichment was assessed by overrepresentation analysis using gseapy (v1.1.12) and the MSigDB_Hallmark_2020 database as previously described (*26*). Lymphocyte-specific cell abundance difference were quantified using Milo as implemented in the pertpy package with default parameters (v1.0.5). In the lymphocyte compartment, mean *Ifng* expression levels were investigated between young and aged mice.

### Generation single nuclear multiomic-sequencing

Both AT2 cells and bronchiolar progenitors were sorted by FACS from two young and two old mice for single-nuclear ATAC and RNA sequencing using a complete sample preparation and library preparation kit (10X Genomics #PN-1000285). In brief, nuclei were isolated from sorted cells with the provided lysis buffer, transposed by Tn5 transposase, GEM-encapsulated, and barcoded using a Chromium Next GEM Chip J (10X Genomics #PN-1000230), following all subsequent manufacturer instructions for library preparation. Proper size distribution of the prepared libraries was confirmed by TapeStation prior to sequencing. Gene expression and ATAC libraries were sequenced separately on an Illumina NovaSeq X Plus platform using an S4-200 flow cell (NovaSeq v1.5 reagent kit) with read configurations of 28+10+10+90 bp for RNA and 50+8+24+49 bp for ATAC (Read 1 + Index 1 + Index 2 + Read 2), at an estimated depth of 70,000 reads per nucleus in both assays.

### Analysis of single nuclear multiomic-sequencing

Raw sequencing data were processed and aligned to a mouse reference genome (mm10) using Cell Ranger ARC v2.0.1 (10x Genomics). Downstream analysis was performed in R v4.2.1 using Seurat v4.3.0 and Signac v1.10.0. Nuclei were filtered using thresholds for RNA (2,000–5,000 detected genes and mitochondrial-to-nuclear gene ratio < 0.10) and ATAC (500–40,000 fragments in peaks, TSS enrichment score > 2, and blacklisted-region fraction < 0.07) separately. Gene expression data were log-normalized and transformed using SCTransform (v0.3.5; 3,000 variable features) with mitochondrial ratio regressed out, followed by principal component analysis. ATAC data were normalized by TF-IDF, dimensionality-reduced by LSI, and scaled with regression of fragment count and nucleosome signal. Both modalities were integrated across all four samples using Harmony v0.1.1 (covariates: sample and age group). Harmony-corrected embeddings were used as input to weighted nearest neighbor (WNN) graph construction (FindMultiModalNeighbors), from which a joint UMAP and SLM-based clusters were derived (FindClusters, algorithm = 3, resolution = 0.2). Cluster marker genes and differentially accessible peaks were identified using FindAllMarkers with Wilcoxon rank-sum (RNA) or logistic regression with fragment count as a latent variable (ATAC). Ambient RNA was detected in non-AT2 cell populations, including many AT2 cell markers, which did not meet the criteria for correction with existing tools (Cellbendr). Therefore, non-AT2 cells were excluded from the analysis to avoid data artifacts. Peaks were subsequently re-called per cell type using MACS2 with ENCODE blacklisted regions excluded, and differential accessibility between young and old mice was assessed within AT2 cells using logistic regression. Using signac, peak-to-gene links in AT2 cells were identified using LinkPeaks and visualized as coverage tracks split by age group using CoveragePlot. To characterize aging transcriptional differences, young and old AT2 cells were subset based on their annotation in the harmonized object and reanalyzed without any integration to visualize the UMAP separation of cells by age in both RNA and ATAC modalities. Differential expression between young and old AT2 cells was assessed using FindMarkers to generate a ranked gene list ordered by mean log2 fold change for Gene set enrichment analysis (GSEA). GSEA was performed using clusterProfiler v4.10.1 with GO Biological Process gene sets (org.Mm.eg.db v3.18.0) and mouse Hallmark gene sets from MSigDB (msigdbr v7.5.1, category = “H”), using a gene set size range of 10–500 and a Benjamini-Hochberg adjusted p-value threshold of 0.05. Computationally intensive work was performed on Harvard University’s O2 high-performance computing cluster.

### Lung organoid culture

Alveolar and airway organoids with supporting stromal cells were largely prepared as before (*3*) with subtle differences in stromal cell isolation. Stomal cells were isolated from 2–4-week-old neonatal mouse lungs (digested with dispase/collagenase as above), including negative selection with Anti-CD31 (Miltenyi Biotech #130-097-418), Anti-CD45 (Miltenyi Biotech #130-052-301), and Anti-EpCAM (Miltenyi Biotech #130-105-958) magnetic beads. Stromal cells were amplified on gelatin-coated plates for 3-5 days before trypsinization and further negative selection to remove contaminating cells. To establish organoids, freshly sorted AT2 cells (1×10^5^ cells/mL) were combined with cultured lung stromal cells (1×10^6^ cells/ml) in a 1:1 mixture of culture media and growth factor reduced Matrigel (Corning #356231). Experiments with bronchiolar progenitors were plated with 2×10^4^ cells/mL or less cells when needed as sorted cells were scarce. Air-liquid interface transwells (Corning #3470) were seeded with 100 uL of this Matrigel mixture and incubated at 37°C for 30 minutes to solidify before adding 500 uL of media on the basolateral side of the transwell. The monoculture of AT2 cells was described previously (*14, 62*). In brief, 2,000 to 2,500 purified AT2 cells were resuspended with 25 μL AT2 Maintenance Medium (AMM) and then mixed with an equal volume of Matrigel (Corning #356255) on ice. A total volume of 50 μL of AMM/Matrigel mixture containing isolated AT2 cells was added to a 96-well plate and AMM was added to the well once the Matrigel was solidified at 37°C for 45 min. The ROCK inhibitor Y-27632 (HY-10583, MCE, USA) was used in the first 4 days of organoid culture and the medium was exchanged every 2-3 days.

### Measurement of organoid formation efficiency and diameter

An overview picture of each well containing organoids was taken using a stereo microscope (SZ×10, Olympus, Japan) 7 days after AT2 cell seeding for feeder-free organoids. The organoid diameter was measured based in these pictures using ImageJ (Version 1.53t). Stroma-supported organoids were counted manually in ImageJ from immunofluorescent images of cultures fixed at day 14, further described below. To calculate the organoid formation efficiency (OFE), we used the number of measured organoids with a size of more than 30 μm: 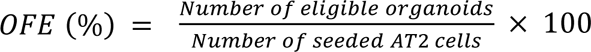.

### Immunofluorescent staining of organoids in matrigel

Organoids in Matrigel-containing transwells were fixed in 10% formalin overnight, washed in PBS, and permeabilized in 0.5% Triton-X/PBS for 1 h at room temperature. Transwells were blocked overnight at 4°C in 10% normal donkey serum/0.2% Triton-X/PBS, then incubated with primary antibodies in blocking solution for two nights at 4°C. Following three washes in Triton-X/PBS, secondary antibodies and DAPI were applied overnight at 4°C in blocking solution, after which transwells were washed three times before imaging. Primary antibodies used were anti-SPC at 1:1000 (Abcam #ab211362), anti-RAGE at 1:100 (R&D #MAB1179-SP), and Anti-Scgb1a1 at 1:100 (Santa Cruz Biotech, sc-390313) with Alexa Fluor-conjugated secondary antibodies applied at 1:100. Transwells were inverted for imaging on a Nikon 90i Eclipse microscope and images were analyzed in ImageJ.

### Immunofluorescent staining of cryosectioned alveolar organoids

At the endpoint of some organoid experiments, the medium was removed prior to the addition of dispase to the wells. The plate was incubated at 37°C for 45 min to allow the Matrigel to dissolve and release the organoids. Organoids were collected, washed with cold PBS twice to remove residual Matrigel, and fixed with 4% PFA, pH 7.0 at RT for 1 h. Fixed organoids were washed with cold PBS twice and subsequently transferred to disposable base molds and embedded with OCT medium (9990426, Thermo Shandon, USA) by freezing them overnight at −80°C. For the preparation of slides, frozen blocks were sectioned in a cryostat (CM3050S, Leica, Germany) with a thickness of 5 μm at the Histology Unit of the Research Center Borstel. All blocks and slides were stored at −80°C until further use. For immunofluorescent (IF) staining of cryosectioned alveolar organoids, slides were placed into a cuvette containing pre-heated citrate acid buffer (10 mM tri-sodium citrate, 0.05% Tween 20, pH 6) in a steamer for 10 min to recover the antigen. Upon antigen retrieval, organoids were permeabilized and blocked with Roti-Block (A151.1, Carl Roth, Germany) supplied with 0.3% Triton X-100 for 1 h at RT. The blocking procedure was followed by primary antibody incubation at 4°C overnight. Primary antibodies were removed and slides were washed with PBST (137 mM NaCl, 2.7 mM KCl, 10 mM Na_2_HPO_4_, 1.8 mM KH_2_PO_4_ and 0.05% Tween-20) three times and then incubated with secondary antibodies (1:1000, goat anti-rabbit, goat anti-rat and goat anti-mouse, all purchased from Thermo Scientific, USA) for 1 h at RT. After washing of the slides with PBST for three times, Mount FluorCare DAPI (HP20.1, Carl Roth, Germany) was used to mount coverslips. Slides were stored in a dark box until imaging at a confocal microscope (SP5, Leica, Germany). We used the following primary antibodies: α-SPC (1:500, AB3786, Millipore), α-RAGE/AGER (1:500, MAB1179-100, R&D System), α-Ki-67 (1:200, 550609, BD Bioscience).

### Lung histology and lung immunofluorescent staining

Following avertin-induced euthanasia mice received a cardiac perfusion of ∼5 mL ice-cold PBS to the right ventricle. All lung lubes or just the left lobe ligated below the primary bronchus with a hemostat were inflated with 4% low-melting point agarose, given a few minutes on ice to solidify, washed in cold PBS wash, and fixed in ice-cold 4% paraformaldehyde with gentle agitation on a rocker overnight at 4°C. Fixed tissue was transferred to 70% ethanol, then processed and embedded in paraffin using standard histological procedures. 5 um lung sections were cut and placed on polarized glass slides before hematoxylin and eosin staining or desiccated for RNA *in situ* hybridization staining described below. H&E staining was analyzed in ImageJ and white balance was corrected with the macro “White balance correction_1.0”. To perform IF staining in paraffin-embedded lungs, slides were subjected to the standard deparaffination process using xylene and ETOH gradients (100%, 95%, 90%, 80%, 70% and 40%). Antigen retrieval of deparaffined lung tissues was conducted by incubating slides with citrate acid buffer (pH 6) in a pressure cooker for 10 min, a microwave oven with 100 Watt for 30 min, or an Epredia Lab Vision PT Module (80400012 Thermo Fisher Scientific). Next, tissues were incubated with Roti-Block supplemented with 0.3% Triton X-100 and 10% goat serum or 0.2% Triton X-100 and 10% donkey serum for 1 h at room temperature. The procedures for antibody staining and coverslip mounting of cyrosectioned organoids described above were used for immunoproteasome KO mice experiments [primary antibodies used: α-SPC (1:500, AB3786, Millipore), α-RAGE/AGER (1:500, MAB1179-100, R&D System), α-PDPN (1: 250, 8.1.1, DSHB). Secondary antibodies were purchased from Thermo Scientific and used at 1:1000 dilutions (goat anti-rabbit, goat anti-rat and goat anti-syrian hamster)].

### RNA in situ Hybridization

mRNA expression and localization in the lung was detected using the RNAscope Multiplex Fluorescent Reagent Kit v2 (Advanced Cell Diagnostics) according to the manufacturer’s protocol (UM323100). Briefly, paraffin-embedded lung sections were baked at 60°C for 1 hour, deparaffinized through two sequential xylene washes (5 min each) followed by two 100% ethanol washes (2 min each), and air-dried. Slides were treated with Hydrogen Peroxide for 10 min at room temperature, followed by antigen retrieval in 1X Target Retrieval Reagent at 98–102°C for 15 min. Protease Plus was applied for 30 min at 40°C in a HybEZ II Oven. Target probes were hybridized for 2 hours at 40°C [Cd3e (314721), Ncr1 (501721-C3), IFNγ (311391-C2), from Advanced Cell Diagnostics], followed by sequential incubation with amplification reagents. HRP-conjugated channel reagents (HRP-C1, HRP-C2, and/or HRP-C3) were applied for 15 min at 40°C, and the fluorescent signal was developed using TSA Vivid dyes (520, 570, or 650, 1:1500 dilution) for 30 min at 40°C. All wash steps were performed in 1X Wash Buffer (2 × 2 min, room temperature). Nuclei were counterstained with DAPI for 30 seconds. After the RNAscope protocol was complete, IF staining was done for AT2 cells with α-SPC (1:1000, Abcam Biochemicals #211326) as described above with minor modifications including a 1:200 dilution of secondary rabbit antibody (Thermo Fisher Scientific #A32790) and mounting with Prolong Gold antifade Mountant (Thermo Fisher Scientific #P10144). Slides were stored at 2–8°C in the dark and imaged within a week on a Nikon 90i Eclipse microscope or Zeiss AxioImager Slide-scanning microscope. Images were analyzed in ImageJ for IFNγ transcript abundance semi-quantitatively by counting the number of TSA vivid signal puncta using the local maxima function. Cd3+ T cell and Ncr1+ NK cell areas were determined by thresholding areas positive for each respective marker signal and puncta numbers were normalized by the area assessed. DAPI+ nuclei within Cd3+ or Ncr+ areas were manually counted as IFNγ+ or IFNγ-based on the presence of any proximate IFNγ puncta.

### Multiplex immunofluorescent staining (mIF)

To determine the cellular composition of the TLS of aged mouse lungs, mIF was employed to stain with multiple antibodies (anti-CD3, -CD8, -NKR, -IFNγ and -PSMB9) in serially sectioned lung slides using the Opal 6-Plex Manual Detection Kit (NEL811001KT, Akoya Bioscience, USA). Paraffin slides were deparaffined as described above. For the antigen retrieval, slides were treated with AR6 Buffer (AR600250ML, Akoya Bioscience, USA) in a microwave oven with 1000W for 1 min and 100W for 10 min. Incubation of the slides with 3% H_2_O_2_ was applied to remove endogenous peroxidase activity. Slides were then washed 3 times with TBST and 1×Antibody Dilute/Block solution (ARD1001EA, Akoya Bioscience, USA) was applied for 10 min at RT to prevent unspecific binding of the antibodies. Subsequently, slides were incubated with diluted primary antibody for 45 min at RT. Upon further washing (3x in TBST), slides were incubated with diluted Opal anti-rabbit HRP-polymer (ARR1001KT, Akoya Biosciences, USA) for 10 min at RT. After washing with TBST, diluted TSA-coupled Opal fluorophore (Akoya Biosciences, USA) was added to the slides for 10 min at RT to amplify the signal. This procedure was repeated for each slide for each antibody and fluorophore. In a final step, diluted Spectral DAPI (FP1490, Akoya Bioscience, USA) was applied to the slides for 5 min at RT prior to coverslip mounting using Prolong Gold antifade reagent (P10144, Thermo Fisher Scientific, USA). All slides were imaged in the same run using a multispectral whole-slide scanner (PhenoImager HT 2.1.0, Akoya Biosciences, USA) at 20x resolution (0.5µm/pixel) using the Motif whole-slide scan with exposure times for all channels kept constant across all slides (DAPI: 1.7 ms, Opal 520: 18.74 ms, Opal 570: 19.76 ms, Opal 620: 41.07 ms, Opal 690: 13.2 ms, Opal 780: 24.07 ms and 100 ms Sample AF). Exposure times were determined and optimized from representative regions across all slides for each target to prevent signal saturation and spill-over into adjacent fluorescence channels. Details for antibody dilution, staining order and used fluorophores are listed in Supplement Table 1.

### Image analysis of mIF

Captured whole-slide images were loaded and processed in Phenochart software (version 2.1.0, Akoya Biosciences, USA) for annotation of the TLS. Images with typical TLS staining from several tissue regions were selected via the inForm software (version 3.0, Akoya Biosciences, USA) for training an algorithm that automatically performed tissue and cell segmentation, as well as cell classification. This algorithm was then applied to all images. The cellular composition of the TLS was calculated based on the percentage of DAPI^+^ cells which express single or multiple markers among total cell number in annotated TLS across all slides. For cell classification, CD3+CD8+ cells were defined as CD8+ T cell and CD3+CD8− cells were defined as CD4+ T cell. We also calculated the number of NK and IFNγ+ cells. Of note, we only included cell-based signals of IFNγ which co-stained for DAPI and IFNγ into our quantification to increase specificity. In total, we analyzed 4 serially sectioned slides for each group.

**Figure S1.**
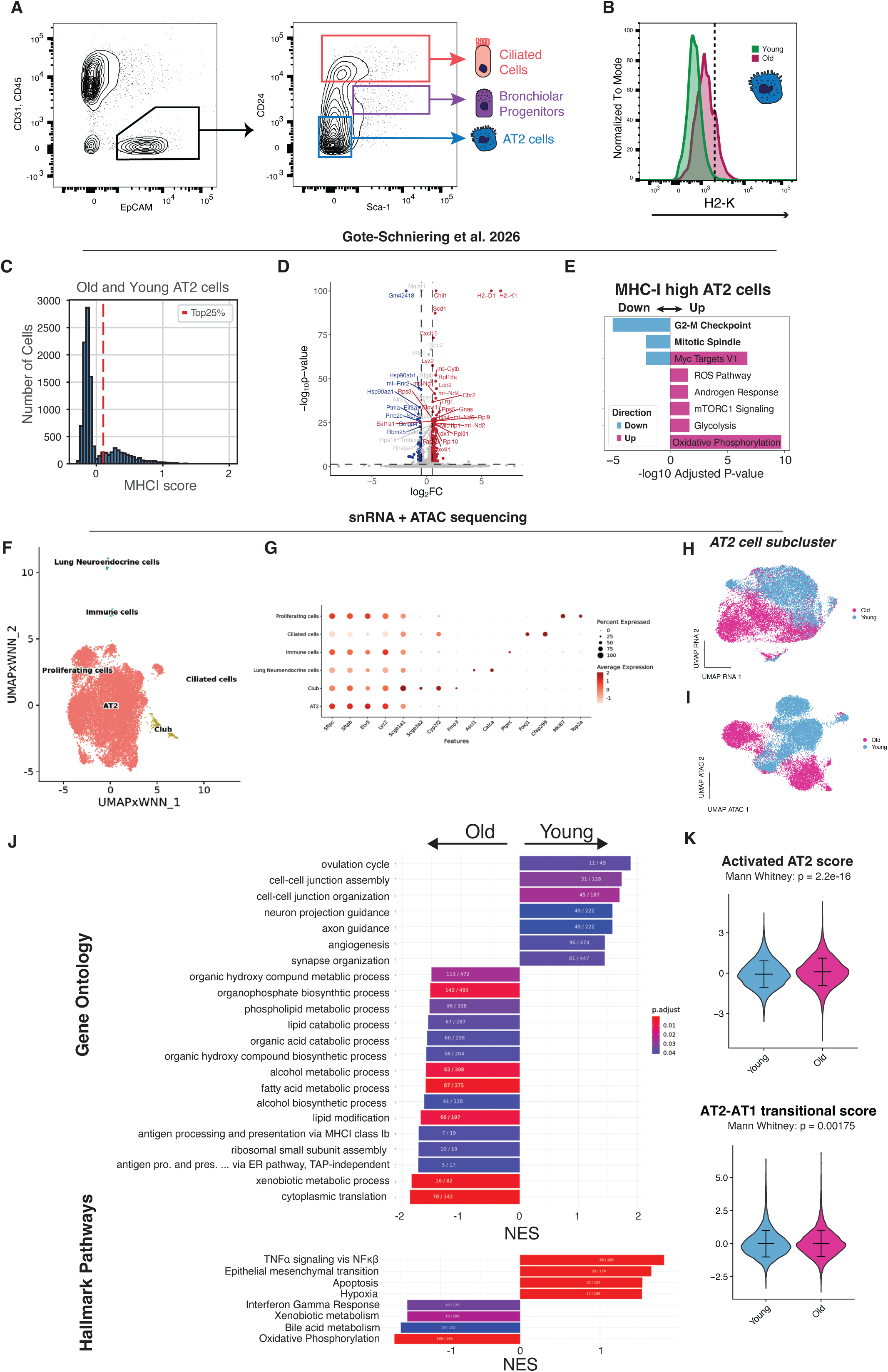
Aged AT2 cells exhibit a conserved inflammatory IFNγ-response signature. A) FACS gating setup to isolate AT2 cells and bronchiolar progenitor cells in old and young mice (*3*). B) Histograms of Anti-H2-K^b^ surface staining during FACS in young and old murine AT2 cells (CD31-CD45-EPCAM+Sca1-CD24-) with dotted line denoting the cutoff for H2-K_b_ hi cells. C) Histogram of MHC-I complex gene score distribution of combined young and aged murine AT2 cells, where the red dotted line indicates the top 25% expressing cells (uninjured lungs from Gote-Schniering et al. *26*). D) Volcano plot displaying the differentially expressed genes in MHC-I complex score high (top 25%) vs rest of AT2 cells. Red color indicates genes that are significantly upregulated with a log2 fold change of >0.5 and adj.p<0.05 and blue color indicates significantly downregulated genes with log 2 fold change < −0.5 and adj.p < 0.05. E) Overrepresentation analysis of differentially expressed genes between the top 25% of AT2 cells by MHC-I complex score and the remaining AT2 cells, where mouse hallmark pathways were assessed (MsigDB). F) A joint UMAP of FACS-isolated AT2 cells and bronchiolar progenitors used for sn-Multiomic sequencing, generated by combining RNA and ATAC assays after harmony integration using a weighted nearest neighbors model. G) Cell-type-specific gene expression used to annotate cell clusters, including elevated AT2 marker genes in non-AT2 cells due to ambient RNA in the single-nuclear pipeline. Independent RNA (H) and ATAC (I) unharmonized UMAPs of AT2 cells show separation of labeled aged cells from young controls. J) GSEA of single nuclear transcriptomic data (Fig 1I) assessed enrichment of MsigDB hallmark pathways and GO pathways based on a ranked gene list. K) Violin plots for transcriptomic signature scores for known AT2 activated and AT2-AT1 transitional states in old and young AT2 cells with published markers (*26*).

**Figure S2.**
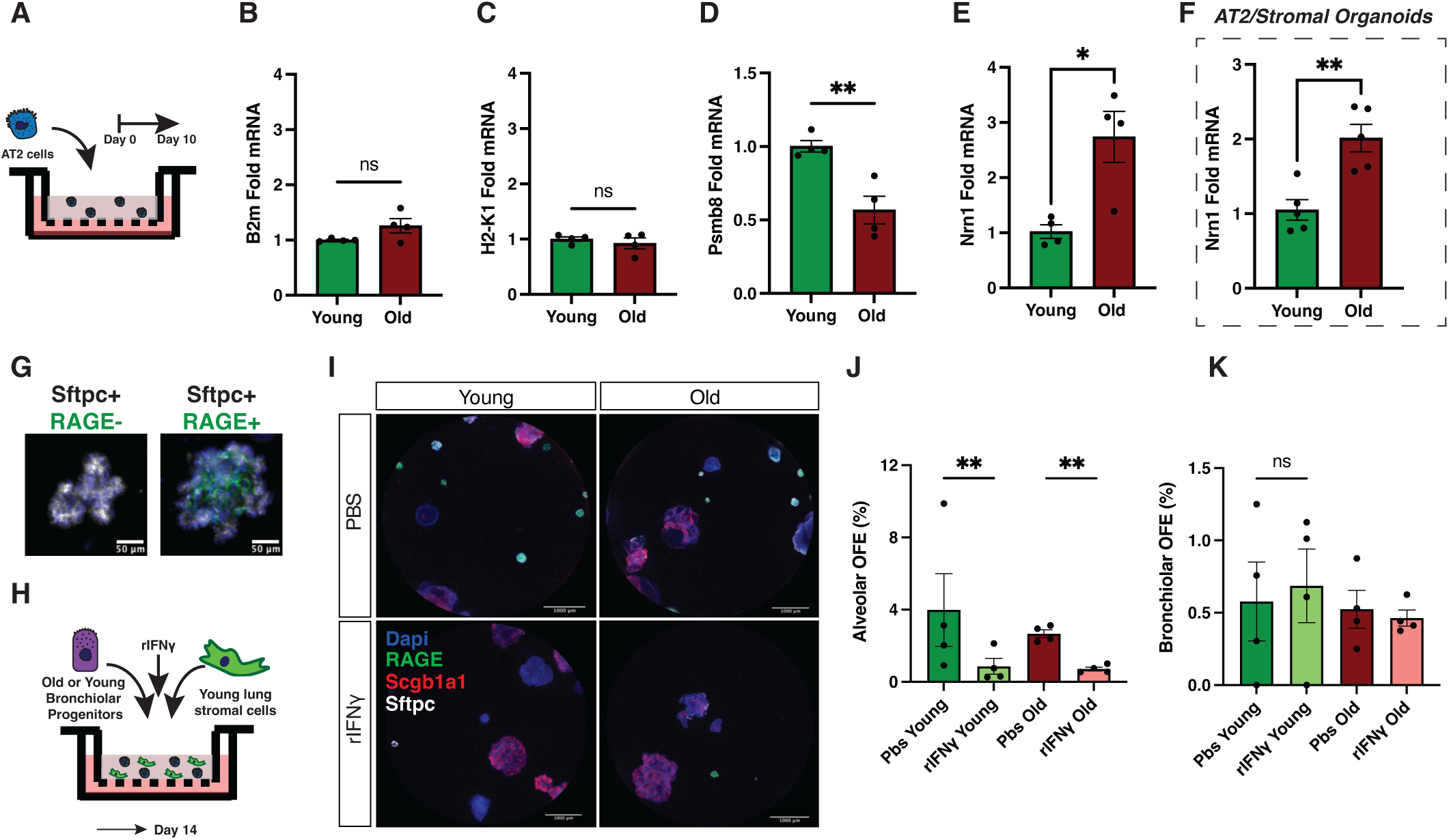
**The IFNγ-response signature of aged AT2 cells is dependent on the aged lung microenvironment and IFNγ disrupts bronchiolar progenitor differentiation to alveolar organoids**. A) qPCR to measure MHCI (B-C) and immunoproteasome(D) gene expression in ex vivo organoid Epcam+ cells at day 10 in feeder-free organoid culture from aged and young AT2. qPCR of aged AT2 chromatin-associated Nrn1 gene expression in (E) feeder-free and (F) stroma-supported organoid cultures from old and young AT2 cells. G) Representative images of alveolar organoids with and without RAGE+ AT1 cell formation, used for Fig. 2D and others. H) Experiment diagram of organoid cultures derived from young or old FACS isolated bronchiolar progenitors after rIFNγ (10ng/mL) or PBS vehicle. I) Representative wells of bronchiolar progenitor-derived organoids with immunofluorescence staining for alveolar (Sftpc and RAGE) and airway cell (Scgb1a1) markers and quantification of organoid efficiency of alveolar (J) and bronchiolar organoids (K). *P < 0.05, **P < 0.01, ***P < 0.001, ****P < 0.0001.

**Figure S3.**
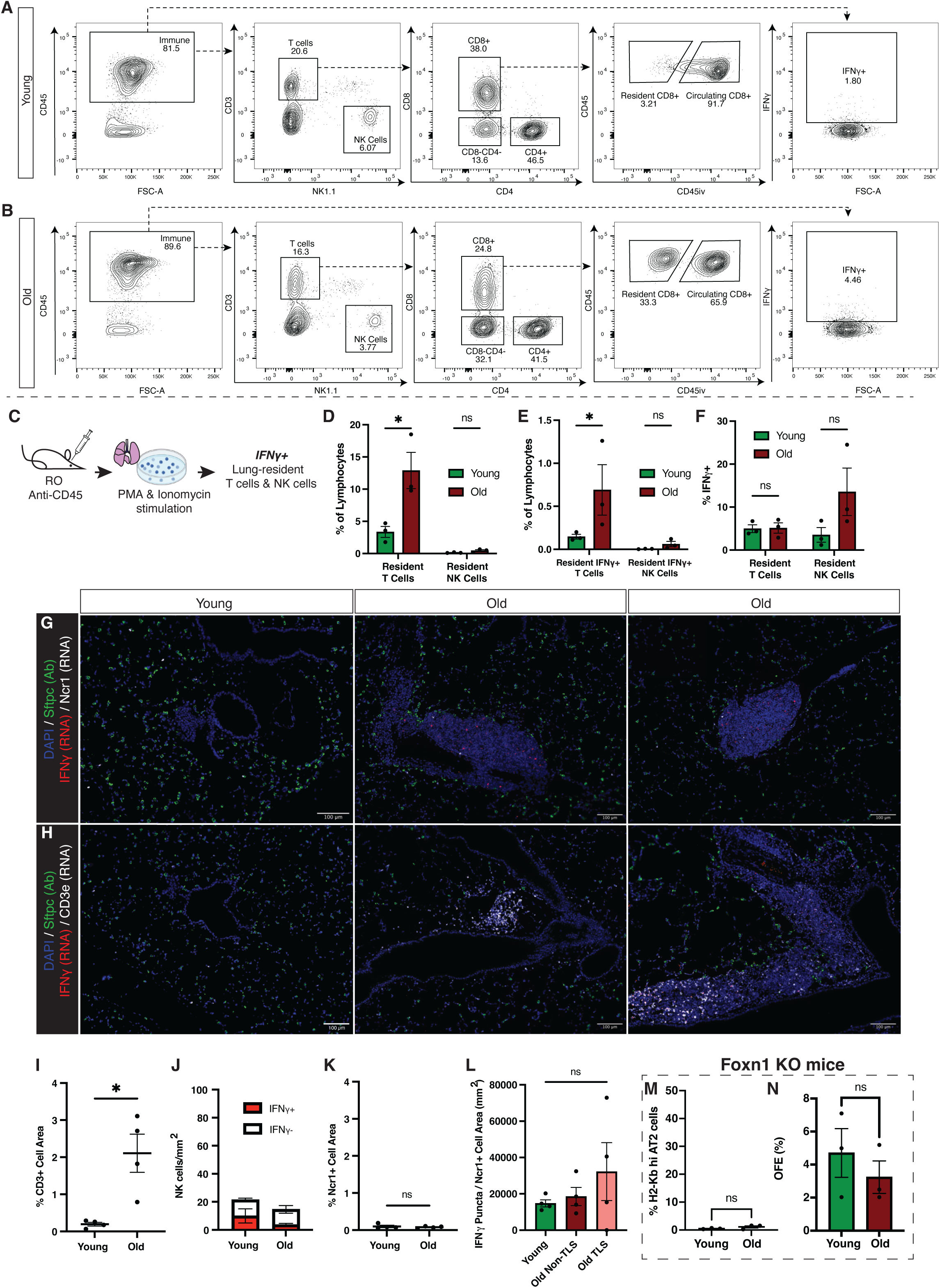
IFNγ+ T cells accumulate within tertiary lymphocyte structures in the aged lung and T cells are required for the aged AT2 cell IFNγ-response signature. A) Representative flow cytometry gating of lung lymphocytes, exclusion of CD45iv+ circulating lymphocytes, and intracellular IFNγ+ cells in either young or old mice (B). C) Experiment diagram of ex vivo lung lymphocytes cultures stimulated to induce IFNγ release followed by flow cytometry of intracellular IFNγ in lung resident lymphocyte populations. D) Subsequent analysis of CD45iv-CD3+ and CD45iv-CD3+ IFNγ+ lung resident T cells (E) as a ratio of all lymphocytes in the old or young mouse lung. F) Qualification of the ratio of IFNγ+ among all resident T cells or NK cells in the old or young lung. G) Representative images of TLS in old lungs or control perivascular areas in young lungs with IF staining of Sftpc (AT2 cells), RNAscope for IFNγ transcript and either Ncr1 transcript labeling NK cells or (H) CD3e transcript labeling T cells. I) Automated Quantification of the ratio of perivascular image area made up of CD3e+ T cells. J) Quantitation of total IFNγ+ Ncr1+ NK cells and (K) Ncr1+ NK cells per mm^2^ in perivascular/peribronchiolar lung regions. L) Quantitation of the density of puncta marking IFNγ mRNA transcript in Ncr1+ cells compared by age and localization within TLS-like structures. M) Quantification of high surface H2-K as a proxy for MHCI complex on AT2 cells by FACS from old (24 months) and young (2 months) Foxn1 knockout mice lacking T cells, as well as (N) organoid-forming efficiency of these AT2 cells in stroma-supported organoid culture. *P < 0.05, **P < 0.01, ***P < 0.001, ****P < 0.0001.

**Figure S4.**
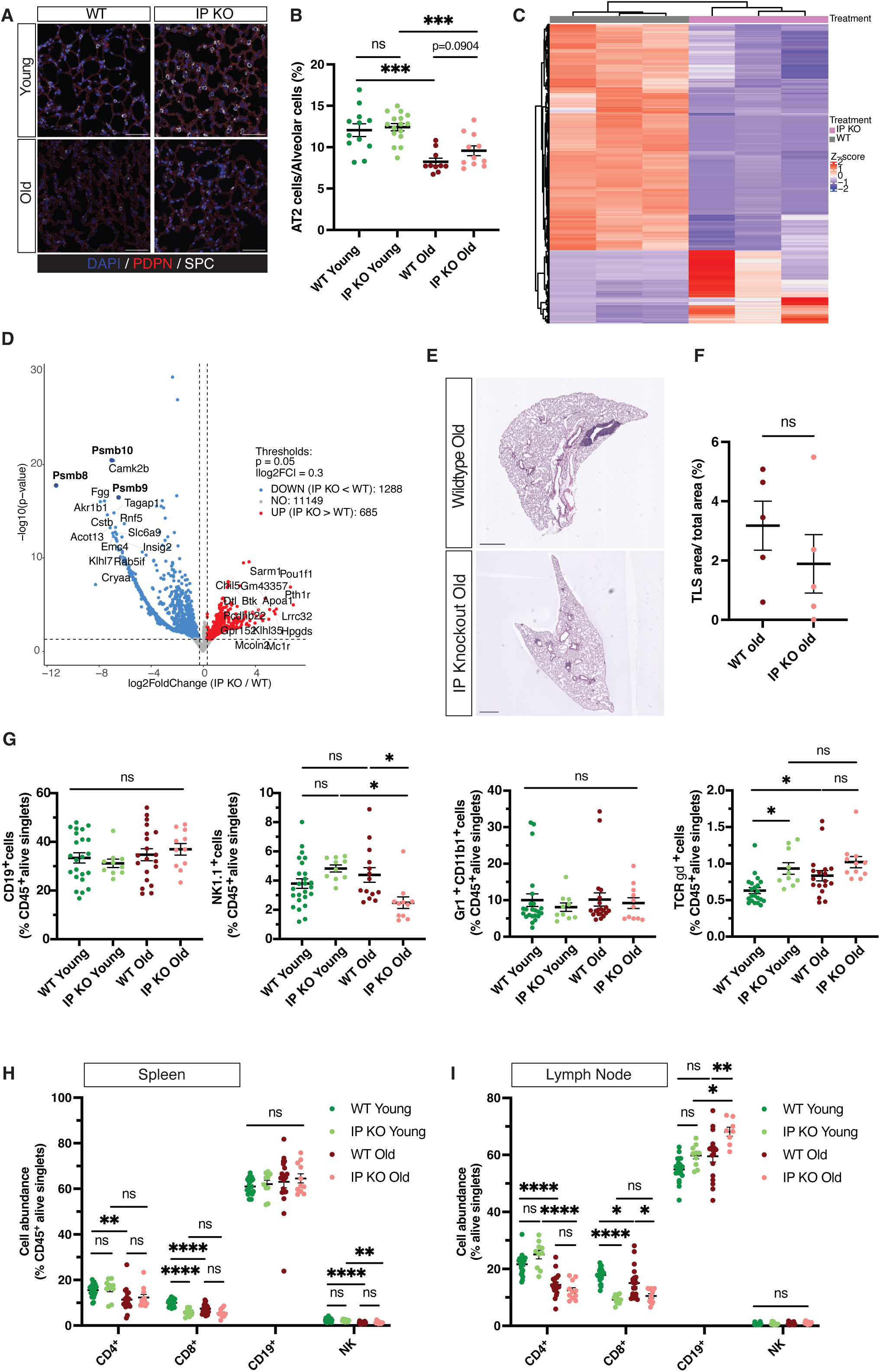
**Deficiency of immunoproteasome attenuates accumulation of CD8+ T cells in aged mice**. A) Representative figure of immunofluorescent staining in young and old lungs from WT or immunoproteasome KO mice for Sftpc and podoplanin (Pdpn). B) Quantification of number of AT2 cell over total alveolar cells within Pdpn+ alveolar regions in young or old lungs from WT or immunoproteasome KO mice. C) Heatmap of differentially expressed genes between immunoproteasome KO and WT groups (padj < 0.05, LogFC <0). D)Volcano plot of differentially expressed genes between immunoproteasome KO and WT groups (padj < 0.05, LogFC <0.3). Top 15 genes in each direction are highlighted. E) H&E staining of aged lungs from WT or immunoproteasome KO mice. F) Proportion of TLS area in total lung area was determined in old WT and immunoproteasome KO mice. G) Indicated immune cell populations shown as % of CD45+ live singlet cells were profiled in lungs of young and old WT or immunoproteasome KO mice using flow cytometry. Immune cell profiles of major immune cell types in (H) spleen and (I) lymph nodes.

**Table S1.**
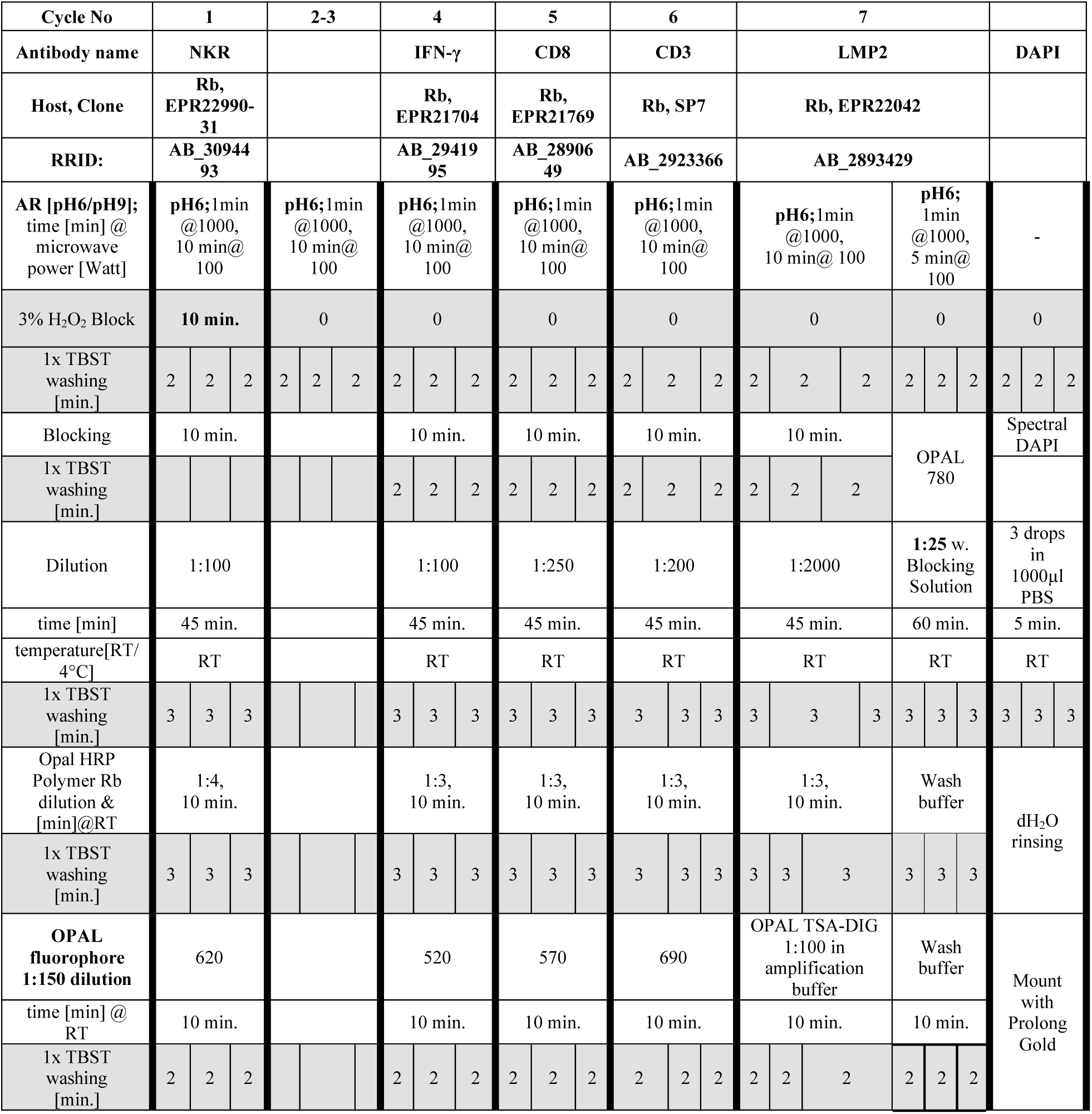
mIF Staining Panel Steps.

